# The rise and fall of the ancient northern pike master sex determining gene

**DOI:** 10.1101/2020.05.31.125336

**Authors:** Qiaowei Pan, Romain Feron, Elodie Jouanno, Hugo Darras, Amaury Herpin, Ben Koop, Eric Rondeau, Frederick W. Goetz, Wesley A. Larson, Louis Bernatchez, Mike Tringali, Stephen S. Curran, Eric Saillant, Gael P.J. Denys, Frank A. von Hippel, Songlin Chen, J. Andrés López, Hugo Verreycken, Konrad Ocalewicz, Rene Guyomard, Camille Eche, Jerome Lluch, Celine Roques, Hongxia Hu, Roger Tabor, Patrick DeHaan, Krista M. Nichols, Laurent Journot, Hugues Parrinello, Christophe Klopp, Elena A. Interesova, Vladimir Trifonov, Manfred Schartl, John Postlethwait, Yann Guiguen

## Abstract

Sexual reproduction is a ubiquitous basic feature of life and genetic sex determination is thus widespread, at least among eukaryotes. Understanding the remarkable diversity of sex determination mechanisms, however, is limited by the paucity of empirical studies. Here, we traced back the evolution of sex determination in an entire clade of vertebrates and uncovered that the northern pike (*Esox lucius*) master sex-determining gene initiated from a 65 to 90 million-year-old gene duplication and remained sex-linked on undifferentiated sex chromosomes for at least 56 million years. Contrasting with its ancient origin, we identified several independent species- or population-specific transitions of sex determination mechanisms in this lineage, including an unexpected complete and recent Y-chromosome loss in some North American northern pike populations. These findings highlight the diversity of the evolutionary fates of master sex-determining genes and raise the importance of careful considerations of population demographic history in sex determination studies. Our study also puts forward the hypothesis that occasional sex reversals and genetic bottlenecks provide a non-adaptive explanation for sex determination transitions.

## Introduction

Genetic sex determination (GSD) evolved independently and repeatedly in diverse taxa, including animals, plants and fungi^1,2^, but the stability of such systems varies drastically among groups. In mammals and birds, the same sex determination (SD) system has been maintained over a long evolutionary time with conserved master sex determining (MSD) genes on sex chromosomes^3–6^. In stark contrast, teleost fish display both GSD and environmental sex determination (ESD)^7,8^, and a remarkable evolutionary lability, driven by dynamic and rapid sex chromosome and MSD gene turnovers^9–12^. These characteristics have made teleosts an attractive model group to study the process of the evolution of sex chromosomes and the turnover of SD systems.

In the past two decades, fueled by advances in genomics tools, a large diversity of MSD genes have been discovered in teleosts, advancing new hypotheses on how the birth of MSD genes, either by allelic diversification or duplication / translocation^9–12^, can drive sex chromosome turnover in vertebrates. Teleost MSD genes also provided empirical support for the “limited option” hypothesis which states that certain genes known to be implicated in the sex differentiation pathways are more likely to be recruited as new MSD genes^9–11,13^. The majority of these recently discovered MSD genes, however, were found in many diverse teleost clades, making it challenging to infer evolutionary patterns and conserved themes in sex chromosomes turnover. Comparative studies on closely related species with SD transitions can thus provide valuable insight on the causes and processes of this turnover^9^. These comparative studies have been done in medakas (*Oryzias sp.*), poeciliids, tilapiine cichlids, salmonids, and sticklebacks, in which different SD systems, sex chromosomes and MSD genes were found even in closely related species^14–18^. Apart from the *Oryzias* family^19–29^ and the salmonids^18,30–32^, however, relatively few studies have explored the evolution of SD systems and the fate of MSD genes within an entire group of closely related species.

Esociformes are a relatively small but old monophyletic teleost order (**Figure 1**) that diverged from a common ancestor around 90 million years ago (Mya) and from their sister clade Salmoniformes about 110 Mya^33,34^. With two families, i.e., Esocidae and Umbridae, and 13 well-recognized species^35^, Esociformes are an ecologically important group of freshwater species from the northern hemisphere^33,34^. Following the identification of a male-specific duplication of the anti-Müllerian hormone, *amhby*, as the MSD gene in northern pike (*Esox lucius*)^36^, we took advantage of the small number of Esociformes species and their relatively long evolutionary history to explore the evolution of sex determination in this entire lineage.

**Figure 1:**
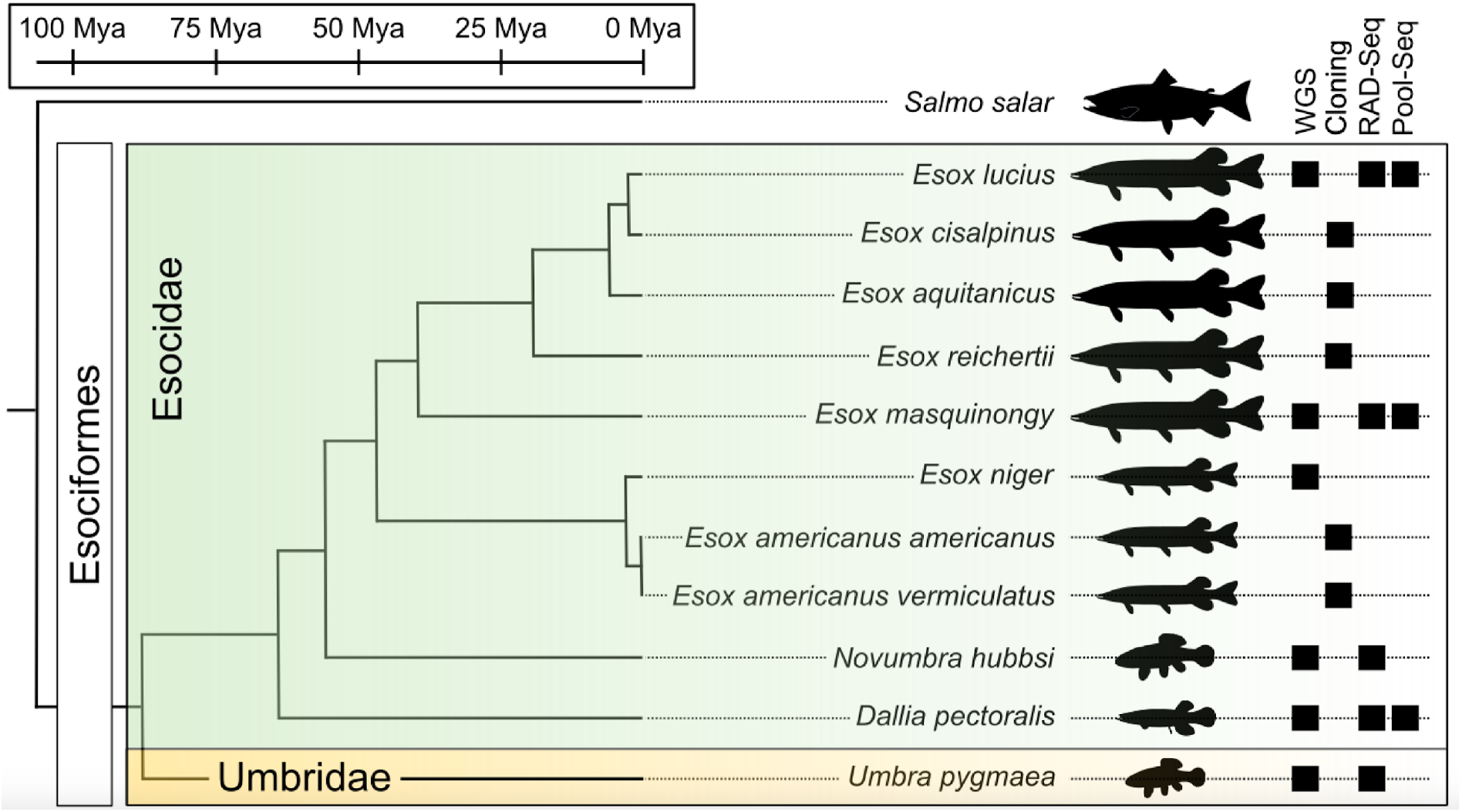
Species phylogeny and technical approaches for investigating sex determination systems in Esociformes. Phylogenetic relationships and estimated divergence time^8,72,73^. The two families within Esociformes, the Esocidae and the Umbridae, are highlighted with green and yellow background, respectively. Whole genome sequencing (WGS), homology cloning (Cloning), Restriction-site associated DNA sequencing (RAD-Seq) and Pooled-Sequencing (Pool-Seq) technical approaches used in each species are shown by a black square on the right side of the figure. Fish silhouettes were obtained from phylopic.org^100^.

Based on whole genome sequencing and population genomic data (**Figure 1**) we traced back the evolutionary trajectory of this MSD gene from its birth in a gene duplication event 65 to 90 Mya to a few independent species- or population-specific SD transitions. Among these SD transitions, we strikingly uncovered even within one species, northern pike, that some populations have experienced a complete and recent loss of their Y-chromosome. Our results highlight the diversity of the evolutionary fates that a single MSD gene can experience in a complete and old monophyletic group of vertebrates. We also show how drift, exacerbated by bottleneck effect, facilitated the loss of a sex chromosome along with its MSD gene in wild populations, providing a first mechanism for such an intriguing entire sex chromosome loss in vertebrates.

## Results

### The *Esox lucius amhby* gene originated in an ancient duplication event

Previously, we identified *amhby*, a male-specific duplicate of the anti-Müllerian hormone gene (*amh)* as the MSD gene in northern Pike (*Esox lucius*). To explore the evolution of *amha* (the autosomal copy of *amh*) and *amhby* in the genus *Esox*, we collected phenotypically sexed samples of most species of Esociformes (**Figure 1** & **Table 1**). Homologs of *amh* were searched by a combination of homology cloning and whole genome sequencing (**Supplementary note 1**) and two *amh* genes were found in all surveyed species with the exceptions of the basally diverging species *Dallia pectoralis* and *Umbra pygmaea* in which only one *amh* gene was found in their whole genome assemblies (**Supplementary note 2**) and in tissue-specific transcriptome databases^1^. To clarify relationships among these *amh* homologs, phylogenetic trees were inferred from these sequences and they provided a clear and consistent topology (**Figure 2A, Figure S1, Supplementary note 3**) indicating an *amh* duplication (*amha* and *amhby* paralogs) that occurred in the common ancestor of the Esocidae^2^. In agreement with that hypothesis, the single *amh* ortholog of *U. pygmaea*, which is the closest sister species to the Esocidae, roots at the base of the *amha* / *amhby* clade. In contrast, the single *amh* ortholog of *D. pectoralis* was systematically placed within the *amha* cluster, strongly suggesting that this species experienced a secondary loss of its *amhby* gene after the duplication of *amh* genes in the Esocidae lineage (**Figure 2B** & **S1**, **Supplementary note 3**).

**Table 1:** Summary of the identification of *amhby* and their association with sex phenotypes in 11 Esociforme species, including six different populations of *E. lucius*, and three different populations of *E. masquinongy*.

| Species | Sampling location | Males with *amhby* | Females with *amhby* | p-value of association with sex |
| --- | --- | --- | --- | --- |
| *Esox lucius* | France | 161/161 (100%) | 60/0 (0%) | **< 2.2E-16** |
| *Esox lucius* | Sweden | 20/20 (100%) | 0/20(0%) | **< 2.2E-16** |
| *Esox lucius* | Poland | 20/20 (100%) | 0/20(0%) | **< 2.2E-16** |
| *Esox lucius* | Xinjiang, China | 5/5 (100%) | 0/5 (0%) | **0.0114** |
| *Esox lucius* | Continental USA and Canada | 0/88 (0%) | 0/74 (0%) | *amhby absent* |
| *Esox lucius* | Alaska, USA | 17/19 (89%) | 0/19 (0%) | **1.79E-07** |
| *Esox aquitanicus* | France | 1/1 (100%) | 0/2 (0%) | 0.665 |
| *Esox cisalpinus* | Italy | 2/2 (100%) | 0/2 (0%) | 0.317 |
| *Esox reichertii* | China | 11/11 (100%) | 0/10 (0%) | **3.40E-05** |
| *Esox masquinongy* | Iowa, USA | 18/27 (66.7%) | 4/18 (22%) | **2.01E-08** |
| *Esox masquinongy* | Quebec, Canada | 9/20 (45%) | 4/20 (20%) | 0.18 |
| *Esox masquinongy* | Wisconsin, USA | 57/61 (93%) | 22/61 (36%) | **1.17E -10** |
| *Esox americanus (americanus)* | Mississippi, USA | 6/6 (100%) | 0/4 (0%) | **0.0123** |
| *Esox niger* | Quebec, Canada | 5/5 (100%) | 0/8 (0%) | **0.00253** |
| *Novumbra hubbsi* | Washington, USA | 21/23 (91%) | 0/22 (0%) | **5.28E-09** |
| *Dallia pectoralis* | Alaska, USA | 0/30 (0%) | 0/30 (0%) | *amhby absent* |
| *Umbra pygmaea* | Belgium | 0/31 (0%) | 0/31 (0%) | *amhby absent* |

**Figure 2:**
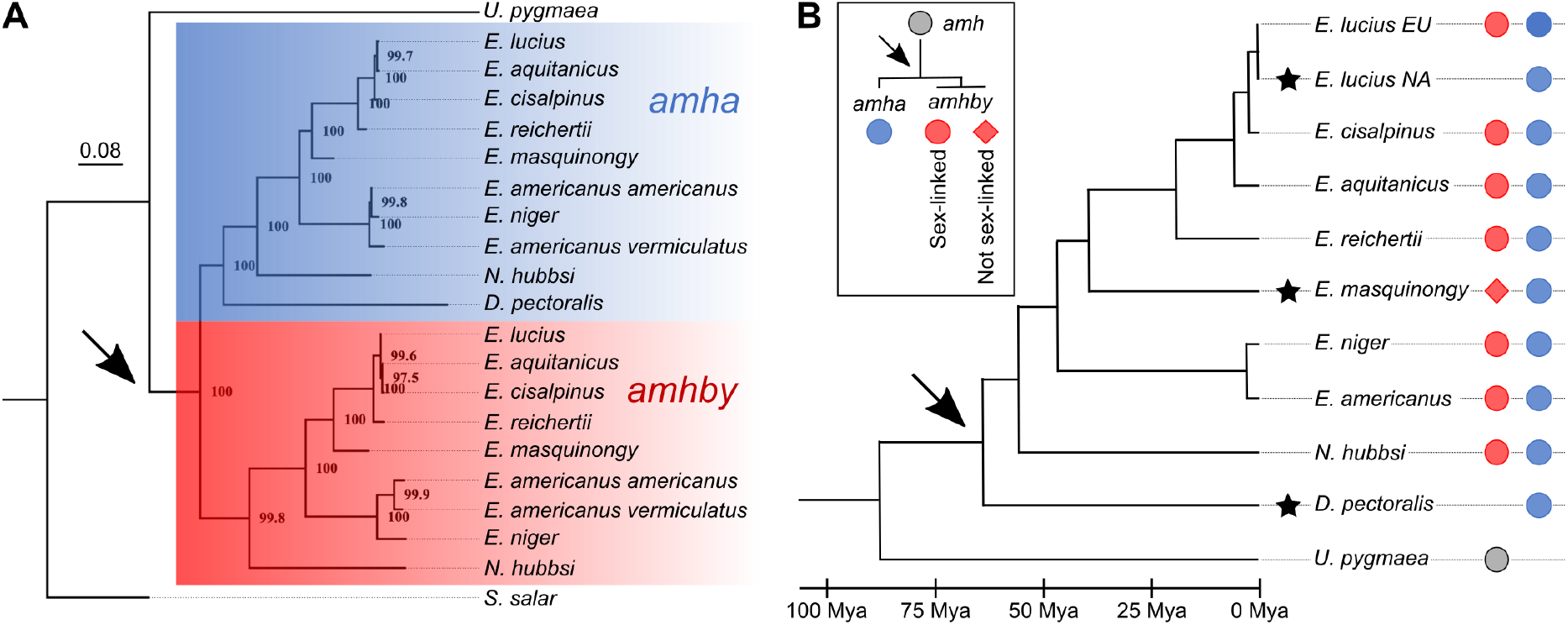
Phylogenetic analysis of *amh* homologs from the Esociformes revealed an ancient origin of *amhby*. **A**: Phylogeny tree of *amh* coding sequences built with the maximum likelihood method. Bootstrap values are given on each node of the tree. The *amha* ortholog cluster is highlighted with blue background and the *amhby* ortholog cluster is highlighted with red background. **B:** Species phylogeny of the Esociformes^8^. The three putative sex determination transitions are shown by a black star. The presence of pre-duplication *amh*, *amha*, and *amhby* along with its sex-linkage is represented by colored dots at the end of each branch. The earliest duplication timing of *amh* is denoted by a black arrow at the root of the Esocidae lineage.

### Sex-linkage of *amhby* in the Esocidae lineage

To explore whether *amhby* is the MSD gene in all these Esocidae species, we first checked if *amhby* is genetically strongly linked to male phenotype, because male sex-linkage would be a strong indication that this gene is located in the non-recombining region of a Y chromosome (**Supplementary note 4)**. Results showed complete sex-linkage in five of the six *Esox* species with the only exception being *E. masquinongy* (**Table 1**), in which significant but incomplete sex-linkage was found in two North American populations from Iowa and Wisconsin, but not in a population from Quebec, Canada (**Table 1**). In addition, heterozygous state of the *amhby* gene was detected in two males and one female of *E. masquinongy* in the population from Quebec (**Figure S2**), conflicting with the expected hemizygous status of a single *amhby* gene on a single Y sex locus. In *Novumbra hubbsi*, the only member of Esocidae not belonging to the *Esox* genus but with an *amhby* gene, we also found a significant but incomplete sex-linkage (**Table 1**).

### Evolution of the structure of the *amhby* gene in the Esocidae

The typical *amh* gene in teleosts comprises seven exons encoding a protein that contains 500 to 571 amino acids depending on the species, with a C-terminal TGF-β domain that is crucial for canonical Amh function (di Clemente et al. 2010; Pfennig et al., 2015). The predicted structures of most of the *amha* and *amhby* genes in Esociformes are consistent with this canonical structure, ruling out an origin by retrotransposition. Furthermore, most *amhby* and *amha* genes do not show any signature of relaxation from purifying selection (**Supplementary note 5**). In both *E. niger* and *N. hubbsi*, however, the predicted Amhby protein is truncated in its N- or C-terminal part. In *E. niger,* this truncated *amhby* gene is flanked by repeated elements and encompasses only three of the seven conserved Amh exons, i.e., exons 5, 6 and an exon 7 with a truncated TGF-β domain (**Supplementary note 6**). In *N. hubbsi*, the truncated *amhby* gene contains the seven conserved Amh exons but with a N-terminal truncation of exon 1 encoding only 8 amino acids with no homology to the conserved 50 amino acid sequences of the first Amh exon of other Esocidae (**Supplementary note 6**). Together, these results show that even in some species having a tight *amhby* sex-linkage, the Amhby deduced protein sequence is strongly modified in its N- and C-terminal conserved domains encoding respectively the TGF-β signal peptide needed for a proper protein secretion and the conserved cysteines of the TGF-β domain required to provide the correct conformation for proper interaction between Amh and its receptor^3^ (**Figure S3**).

### Whole genome analyses of the evolution of sex-determination systems in Esociformes

To complement *amhby* sex-linkage analyses, we also searched for whole-genome sex-specific signatures in species with either incomplete *amhby* sex linkage (*E. masquinongy* and *N. hubbsi*) or species without *amhby* (*D. pectoralis* and *U. pygmaea*). For species with incomplete linkage of sex to *amhby*, we produced gDNA Pool-sequencing (Pool-Seq) of *amhby* positive males versus *amhby* negative females in the Iowa population of *E. masquinongy* and gDNA RAD-sequencing (RAD-Seq) of phenotypic males versus phenotypic females in *N. hubbsi*. Results from these analyses (**Supplementary notes 7 and 8**) provided for these two species a low level of male-specific signal that indicates a small sex locus region with a male heterogametic SD system (XX/XY) (**Figure 3A & Figure S5**). For species without *amhby*, RAD-Seq of phenotypic males versus phenotypic females was carried out in both *D. pectoralis* and *U. pygmaea* (**Supplementary note 7**). In *D. pectoralis,* only three female-biased RAD markers were identified, suggesting that this species has a small sex locus region under a female heterogametic SD system (ZZ/ZW) (**Figure 3B**). This ZZ/ZW SD system was further supported by a substantially higher number of female-specific sequences in Pool-Seq analysis (**Supplementary note 7)**. In contrast, a large number (N = 137) of male-biased RAD markers were identified in *U. pygmaea,* that does not have the *amhby* gene, supporting the hypothesis that this species has a large sex locus region under a male heterogametic SD system (XX/XY) (**Figure 3C**).

**Figure 3:**
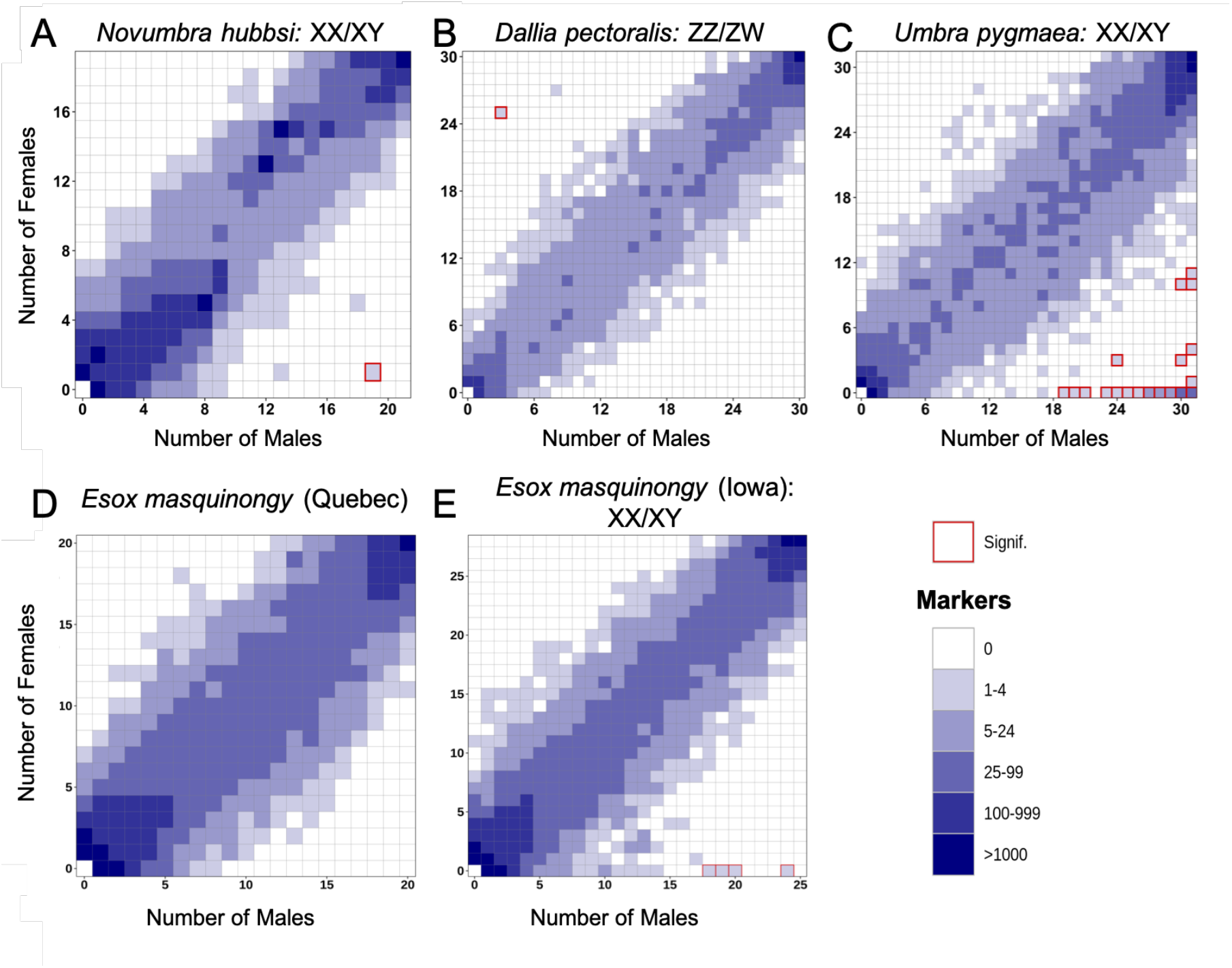
Characterization of *s*ex determination systems in different Esociformes species through RAD-Seq analyses. Each tile plot shows the distribution of non-polymorphic RAD-Seq markers shared between phenotypic males (horizontal axis) and phenotypic females (vertical axis). The intensity of color for a tile reflects the number of markers present in the respective number of males and females. Tiles that are significantly associated with phenotypic sex (Chi-squared test, p<0.05 after Bonferroni correction) are highlighted with a red border. **A**: In *N. hubbsi*, two markers (indicated by the shade corresponding to 1-4 markers) were present in 19 males but only one female, indicating significant male association; reciprocally, no marker was present in many females but few males. This result indicates an XX/XY sex determination system with a small sex locus. **B**: In *D. pectoralis*, one marker showed a significant association with females while no marker showed association with males, indicating a ZZ/ZW sex determination system. **C**: In *U. pygmaea*, 140 markers showed a significant association with males, while no marker showed association with females, indicating a XX/XY sex determination system with a large non-recombining sex locus. **D**: In the population of *E. masquinongy* from Quebec (Canada), no marker was found to be associated with either sex. **E**: In a population of *E. masquinongy* from Iowa (USA), analysis identified five markers significantly associated with male phenotype, indicating a XX/XY sex determining system.

### Some populations of *Esox lucius* lost their Y chromosome and ancestral master sex determining gene

Although *amhby* was demonstrated to be the MSD gene in European populations of northern pike^4^, this gene was surprisingly absent from the high quality genome assembly of a male specimen from a Canadian population (GCA_000721915.3)^5^. To better explore this discrepancy, we characterized the *amhby* sex linkage in several geographically isolated populations of *E. lucius* and found significant male sex linkage in all investigated populations from China, Europe and Alaska (**Table 1**, **Supplementary note 9**). In contrast, *amhby* was found to be completely absent, both in males and females, from all Canadian and mainland USA populations (**Table 1**).

To investigate whether the loss of *amhby* coincides with large genomic changes, we first compared phenotypic males and phenotypic females from a European population carrying the *amhby* gene (Ille-et-Vilaine, France) with a North American population that lost its *amhby* gene (Quebec, Canada) using a Pool-Seq approach. These Pool-Seq results were aligned to an improved European *E. lucius* genome assembly (NCBI accession number SDAW00000000) in which all previously identified Y-specific contigs^4^ were scaffolded into a single contiguous sex locus on the Y (LG24) sex chromosome (**Table S1**). In the European population of *E. lucius*, Pool-Seq analysis (**Supplementary note 9**) confirmed, but with a much better resolution, previous results^4^, showing that this European *E. lucius* population harbors a rather small (140 kb) non-recombining sex locus at the proximal end of LG24 (**Figure 4A.1 & 4B.1**). In the Canadian population, virtually no reads from either the male or female pools mapped to this 140 kb Y-specific region identified in the European population (**Figure 4A.2 & 4B.2**) and no sign of differentiation between males and females could be observed along the rest of LG24. In the Quebec population, not only LG24 but the entire rest of the genome had no sex-specific heterozygosity for either male or female reads with either the Pool-Seq analysis or a reference-free RAD-Seq approach (**Figure** S**5, Supplementary note 9**). Together, these results suggest that this Canadian population, and likely other mainland USA populations lack the male-specific Y region found in European populations, including the MSD gene *amhby.* Moreover, these North American populations did not acquire a sufficiently large new sex locus that could be detected by the RAD-Seq and Pool-Seq approaches, or their current SD relies on a non-genetic mechanism.

**Figure 4:**
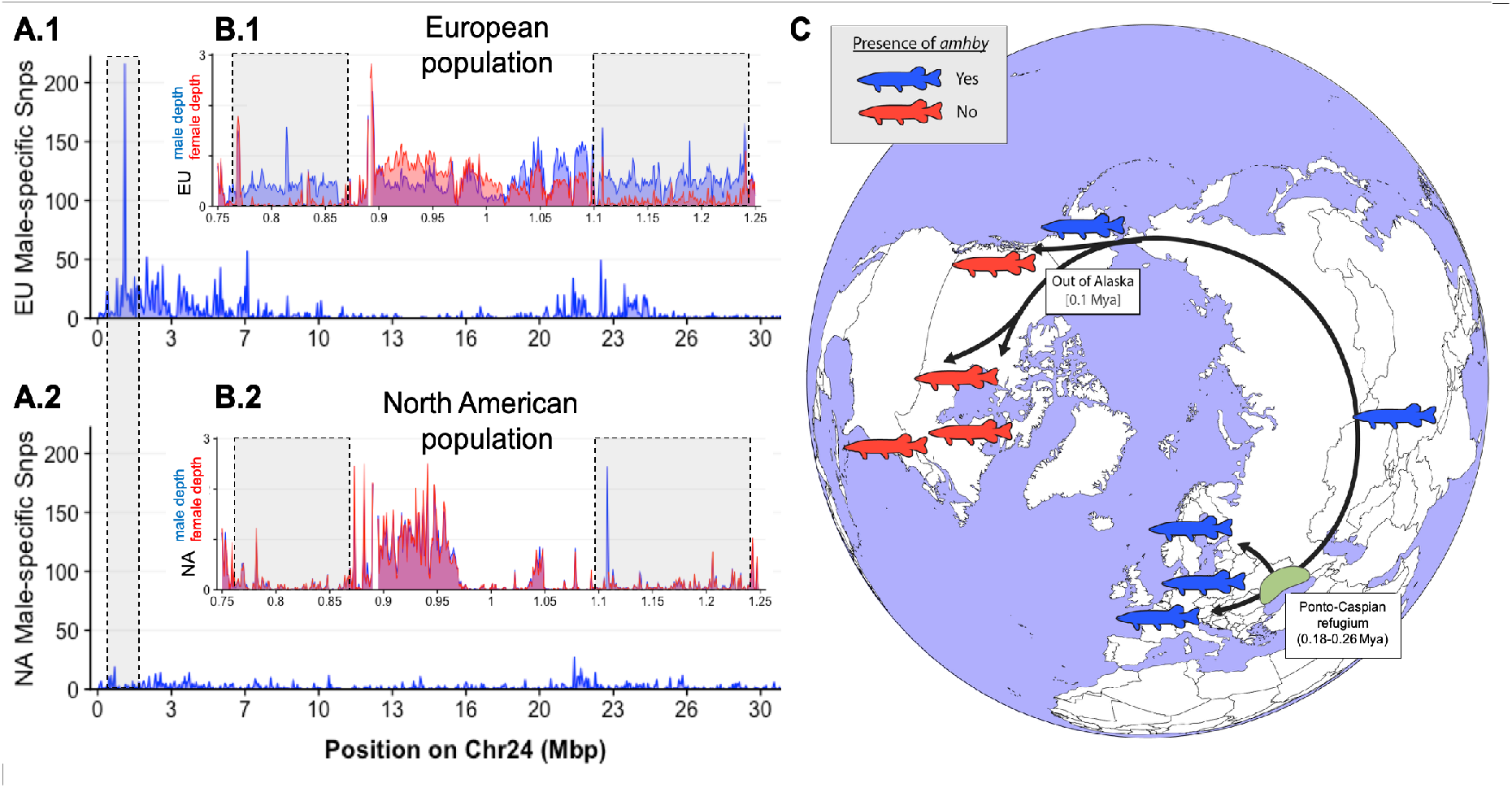
Complete loss of the ancestral sex locus and master sex determination gene in North American Northern pike (*Esox lucius*). **A**: The numbers of male- and female-specific SNPs in 50 kb non-overlapping windows deduced from Pool-Seq data of a European population (**A.1**) and North American population (**A.2**) of *E. lucius* are plotted along the Y chromosome (LG24) of the sequence of *E. lucius* from a European male genome (SDAW00000000). Male data are represented in blue and female in red. While the highest number of male specific SNPs is found in a single window located at the proximal end of LG24 (0.95 Mb to 1.0 Mb) that contained 182 male-specific SNPs (highlighted by the gray box) in the European *E. lucius* population, the same region showed no differentiation between males and females in the North American population. **B:** Relative coverage of male and female Pool-Seq reads is indicated by blue and red lines, respectively, in contigs containing *amhby* in the European population (**B.1**) and in the North American population (**B.2**). While in the European population, this region is only covered by male reads at a coverage depth close to 0.5 genome average, it is covered by no male nor female reads in the North American population, indicating the loss of the entire Y-specific region, highlighted by the gray boxes. **C:** Schematic representation of one hypothesized route of post-glacial *E. lucius* expansion from a Eurasian refugia ~ 0.18 to 0.26 Mya and the presence (red silhouettes) / absence (blue silhouettes) of *amhby* in different populations, showing that this master sex determining gene was lost in the North American population during this out-of-Alaska dispersal around 0.1 Mya. The hypothesized refugium in the Ponto Caspian region is indicated with a yellow highlight. An alternative route of post-glacial dispersal is shown in **Figure S6**.

### Evolving sex determination systems in the muskellunge, *Esox masquinongy*

As shown above, variable *amhby* sex-linkages were observed among different populations of *E. masquinongy* (**Table 1**). We then searched genome-wide for sex-specific signals using RAD-Seq in a population from Iowa (US) in which *amhby* was incompletely but significantly associated with male phenotype and in a population from Quebec (Canada) where *amhby* association was not significant. In the Iowa population, five RAD markers were significantly associated with male phenotype, while no marker was associated with female phenotype (**Figure 3D**); this result indicates a male heterogametic sex determination system (XX/XY) with a low differentiation between the X and Y chromosomes as observed in the northern pike^4^ and *N. hubbsi* (this study). In contrast, no sex-specific marker was identified in some Canadian populations (**Figure 3E, Figure S7, Supplementary note 7)**.

## Discussion

Previous results on the *amhby* MSD gene of northern pike^4^ showed that its sequence was substantially divergent from its autosomal counterpart *amha*, in contrast to other known cases of *amh* duplication in fishes^6,7^, suggesting that this MSD gene may have an ancient origin and / or may have undergone an accelerated evolution. Here, we show that the duplication that produced *amhby* occurred before the last common ancestor of Esocidae in all species of *Esox* and *Novumbra*, but was subsequently lost in the *Dallia* lineage. Esocidae diverged from the Umbridae between 65 to 90 Mya^8^, hence the high level of differentiation observed between *amha* and *amhby* is likely due to the ancient origin of the duplication event. We also demonstrate that *amhby* is completely sex-linked in the majority of Esocidae species in which this gene was retained with low levels of sex chromosome differentiation at least in some populations of *E. masquinongy* and *N. hubbsi*. Taken together, these results suggest that *amhby* likely acquired an MSD function, either 56 Mya in the last common ancestor of *Esox* and *Novumbra*, or 66 Mya in the common ancestor of all Esocidae followed by a subsequent loss in *D. pectoralis*.

This SD system has been maintained for more than 56 My without driving the evolution of highly differentiated sex chromosomes, similar to what was previously described in *E. lucius*^4^. The identification of an old MSD gene located on a homomorphic sex chromosome with limited differentiation over a long evolutionary time is thus in stark contrast with classic models of sex chromosome evolution that postulate the gradual decay of sex chromosomes^9^. Previous work on cichlid fishes and true frogs suggested that frequent turnovers could keep sex chromosomes undifferentiated^10,11^. It was proposed for these organisms that autosomes that are already hosting genes involved in sex differentiation pathways could be recruited as new sex chromosomes, facilitating turn overs of sex chromosomes and MSD genes^12^. Alternative evolutionary pathways could also involve conserved MSD genes that translocate onto different autosomes as has been found for the salmonid *sdY* gene^13,14^. In addition, frequent sex reversal of the heterogametic sex has been proposed to ‘rejuvernate’ sex chromosomes and keep them homomorphic^15^. Our results, based on whole genome analysis in *E. masquinongy*, suggest that the same small Y chromosome sex locus has been conserved in most Esocidae. Such a stability is unusual in teleosts, where frequent turnover of SD systems is considered the norm^16–19^. In line with our present results, however, substantially undifferentiated sex chromosomes have also been found to be maintained over relatively long evolutionary periods in some Takifugu fish species^20^, supporting the idea that theoretical models cannot be generalized and that additional and as yet unknown evolutionary forces can prevent important sex chromosome decay.

Even though *amhby* is an ancient and conserved MSD gene, this old MSD gene experienced different fates across the Esocidae lineage. One of these unexpected fates is the evolution of the Amhby protein in *N. hubbsi*, and *E. niger*, which lack parts of the conserved C- and /or N-terminal Amh regions^21^. In these two species, *amhby* was significantly and almost completely sex-linked, suggesting that this gene could still function as a MSD gene. Because the conserved Amh domains that are altered in the *amhby* genes in these two species are needed for proper protein secretion and interaction between Amh and its receptor^3^, it is unclear how these truncated Amhby proteins could still serve as functional MSD proteins. A similar duplication and truncation of an *amh* MSD gene has already been reported in the cobaltcap silverside (*Hypoatherina tsurugae*), another teleost^22^. In addition, other cases of MSD genes having evolved through duplication / truncation of their ancestral copy have also been described, such as the *sdY* MSD gene in Salmonids and the putative *amhr2by* MSD gene in yellow perch, *Perca flavescens*, demonstrating that preservation of all ancestral domains is not always necessary for a duplicated protein to assume an MSD role^13,23,24^. Because domain gains and losses can both contribute to new protein functions^25,26^, the evolution of a new MSD protein structure can even be seen as a strong evolutionary driver required for the emergence and fixation of some new MSD genes.

Another striking evolutionary fate of this ancient *amhby* SD system in Esocidae relies on the several independent SD transitions that we characterized in this lineage. The first is a complete transition in *D. pectoralis* from an XX/XY male heterogametic SD system, which is shared by Esocidae and *U. pygmaea*, to a ZZ/ZW female heterogametic SD system. Interestingly, this SD transition was also accompanied by, or shortly followed by, a secondary loss of *amhby* because our phylogenetic analyses consistently support the timing of the *amhby* duplication in the common ancestor of the Esocidae i.e., before the divergence of the *D. pectoralis* lineage. While all other Esociformes uniformly display a clear XX/XY SD system, this ZZ/ZW SD system in *D. pectoralis*, could suggest that the shift from male to female heterogamety was potentially driven by the loss of the *amhby* MSD gene, because this kind of SD transition has been documented in other species, for example, in the housefly^27^. The other SD transitions that we characterized, those in *E. lucius* and *E. masquinongy,* are probably more recent because they appear within populations of the same species and not between different species. In *E. masquinongy*, evidence from whole genome analysis and *amhby* sex-linkage both support its conservation as a MSD gene along with a clear XX/XY SD system in two USA populations. In contrast, in a Canadian population from Quebec, our whole genome analysis did not support a male heterogametic SD system. In addition, despite being present in this population, *amhby* was no longer sex-linked and a few males and females even displayed allelic variations in the *amhby* gene, which is not compatible with the expected hemizygous status of a Y sex locus gene. This finding could suggest that the *amhby* locus in this population is not in the non-recombining sex region and thus is no longer an MSD gene. Because we did not find *amhby* sequence differences among populations of *E. masquinongy*, this result would suggest that this SD transition is linked to the recent emergence of a new population-specific SD mechanism. The fact that SD was suggested to be female heterogametic in some populations of *masquinongy*^28^, and that the great lakes population and the Quebec population belong to different lineages^29^, further support the hypothesis of population-specific variation in *E. masquinongy* SD. But an interesting alternative hypothesis would be that this SD system relies on a simple monofactorial XX/XY system in some *E. masquinongy* populations and on a polygenic system with multiple sex chromosomes in other populations of the same species. This hypothesis would equally fit with our current results because it would explain both the absence of complete sex-linkage and the heterozygosity of *amhby* in some populations. This explanation would also fit with the suggested female heterogamety of some populations, which was inferred from the presence of males in the offspring of gynogenetic females^28^, a result that could equally be the indication of a ZZ/ZW monofactorial system or of a polygenic system with multiple sex chromosomes. Such polygenic SD systems, including systems with multiple sex chromosomes^30,31^, are complicated to characterize but they have been well described in a limited number of species^30^ and even in some populations of the same species such as the Poeciliids *Xiphophorus nigrensis* and *X. multilineatus*^32^.

In *E. lucius*, we found that *amhby* is present and completely sex-linked in all populations we surveyed from Europe, Alaska and Asia, suggesting that this gene functions as an MSD gene in all of these populations. The entire Y sex locus including *amhby,* however, was absent in some North American populations, suggesting that these populations have lost their ancestral Y sex chromosome and that their remaining X chromosomes reverted to autosomes. The current geographic distribution of *E. lucius* results from a post-glacial expansion from three glacial refugia ~ 0.18 to 0.26 Mya, and the North American populations without *amhby* belong to a monophyletic lineage with a circumpolar distribution originating from the same Eurasian origin^33^. Fossil records from Alaska support the idea of an “out of Alaska” North-American expansion of *E. lucius* within the last 100,000 years^34^, suggesting that the *amhby* sex locus could have been lost during this dispersal period (**Figure 4C**). The complete loss of this sex locus in such a short evolutionary time is unlikely to result from the slow pseudogenization of the MSD gene as was found for instance in the Luzon medaka, *Oryzias luzonensis*^35^. Rather, it is probably the result of the loss of the entire Y chromosome in founder individuals. This founder hypothesis is supported by the fact that North American populations of *E. lucius* likely experienced a strong bottleneck because they display a much lower genetic diversity compared to other circumpolar lineage populations^33^. The loss of the Y chromosome could then stem from founder males being genetic XX females that experienced sex reversal; spontaneously masculinized XX fish, likely due to environmental effects, are occasionally observed in captive *lucius*^4^. Environmental influence on GSD is a well documented phenomenon found in many fish species, such as Nile tilapia^36^, Half-smooth tongue sole^37^ and Japanese rice fish^38^. These environmental effects on SD may result in the persistence of populations which would otherwise go extinct following the loss of the ancient MSD gene. Notably, this Y chromosome loss in natural populations of *E. lucius* mirrors the loss of the W chromosome in lab strains of zebrafish after just a few decades of selective breeding^39^, with the current SD mechanism in laboratory stocks hypothesized to be polygenic^40^, environmental^41^ or even ‘random’^42^, having evolved from the wild, natural ZW strains. Whether ESD, which is common among poikilothermic animals including teleosts^43^, could be the only SD system in these North American populations or whether a new MSD gene has taken the lead of a new GSD system is still unresolved, but our whole-genome analyses failed to identify any sex-associated markers in these North American populations, meaning that if such a new sex locus exists, it would lack detectable signatures of molecular differentiation like what has been found for instance in some *Takifugu* species^20^.

A few evolutionary models have been formulated to capture the dynamics of sex chromosome turnover^12^. The replacement of the ancestral SD locus by a new one could occur via neutral processes driven by drift alone, or via positive selection when the new MSD gene confers a fitness advantage^44^, or when this new MSD gene is associated with strong sexually antagonistic alleles^45^. Moreover, the accumulation of deleterious mutations on the non-recombining sex locus of the sex chromosome could facilitate sex chromosome turnover^46^. In all these models, the ancestral sex chromosomes would revert to autosomes only when the new SD locus is fixed in the population. In North American populations of *E. lucius*, the ancestral sex chromosome was likely lost due to sudden drift in bottlenecked populations. Interestingly, this process would not necessarily need the simultaneous emergence of a new GSD system, given the flexibility of SD mechanisms in teleosts, in which environmental cues can compensate at least temporarily for the absence of a strict GSD system. ESD, which is known to be easily invaded by new MSD^47,48^, could then serve as a transitional state in between sex chromosome turnovers.

Collectively, our results depict the evolutionary trajectories of a conserved MSD gene in an entire clade of vertebrates, highlighting both the potential stability of MSD genes as well as non-differentiated sex chromosomes in some lineages, and also the dynamics of species- or population-specific independent SD evolution events in teleost fish. Our results also point to the importance of careful consideration of the population demographic history of SD systems, and of taking into account the potential buffering role of ESD during transitions of SD systems in models of sex chromosome evolution.

## Methods

### Sample collection

Information on the different Esociform species, including species collectors, sexing method and experiments performed is given in **Table S2**.

### Genomic DNA extraction

Fin clips were collected and stored at 4°C in 75-100% ethanol until genomic DNA (gDNA) extraction For genotyping, samples were lysed with 5% Chelex and 25 mg of Proteinase K at 55°C for 2 hours, followed by incubation at 99°C for 10 minutes^49^. For Illumina sequencing, gDNA was obtained using NucleoSpin Kits for Tissue (Macherey-Nagel, Duren, Germany) following the producer’s protocol. DNA concentration was quantified using Qubit dsDNA HS Assay Kit (Invitrogen, Carlsbad, CA) and a Qubit3 fluorometer (Invitrogen, Carlsbad, CA). For Pool-Seq, DNA from different samples was normalized to the same quantity before pooling for male and female libraries separately. High molecular weight gDNA for long read sequencing was extracted as described by Pan et al. (2019)^4^.

### Genome and population genomics sequencing

Draft genomes of northern pike (*Esox lucius*), muskellunge (*Esox masquinongy*), chain pickerel (*Esox niger*), Olympic mudminnow (*Novumbra hubbsi*), and Alaska blackfish (*Dallia pectoralis*) were sequenced using a whole genome shotgun strategy with 2×250 bp Illumina reads. Libraries were built using the Truseq nano kit (Illumina, ref. FC-121-4001) following instructions from the manufacturer. 200 ng of gDNA was briefly sonicated using a Bioruptor (Diagenode). The gDNA was end-repaired and size-selected on beads to retain fragments of size around 550 bp, and these fragments were a-tailed and ligated to Illumina’s adapter. The ligated gDNA was then subjected to 8 PCR cycles. Libraries were checked on a Fragment Analyzer (AATI) and quantified by qPCR using a library quantification kit from KAPA. Libraries were sequenced on a HIseq2500 using a paired end 2×250 bp v2 rapid mode according to the manufacturer’s instructions. Image analysis was performed by HiSeq Control Software and base calling by the RTA software provided by Illumina. The output of each run can be found in **Table S3**. For improving our European male genome of *E. lucius*, we generated an extra coverage of Oxford Nanopore long reads using a higher fragment size (50 kb) library made from gDNA extracted from a different male from the same European population as the sample used for previous genome assembly. Library construction and genome sequencing was carried out as previously described^4^ and 12.7 Gbp of new data were generated from one PromethION flowcell.

Pool-Seq was performed on the North American population of *E. lucius*, on *E. masquinongy*, and on *D. pectoralis*. Pooled libraries were constructed using a Truseq nano kit (Illumina, ref. FC-121-4001) following the manufacturer’s instructions. Two sex-specific DNA Pool-Seq libraries were prepared for each species using the Illumina TruSeq Nano DNA HT Library Prep Kit (Illumina, San Diego, CA) with the same protocol as for the draft genome sequencing. The libraries were then sequenced on a NovaSeq S4 lane (Illumina, San Diego, CA) using paired-end 2×150 bp mode with Illumina NovaSeq Reagent Kits following the manufacturer’s instructions. The output of each run for each sex can be found in **Table S3**.

RAD-Seq was performed on the North American population of *E. lucius*, on *E. masquinongy*, on *N. hubbsi*, on *D. pectoralis* and *U. pygmaea*. RAD libraries were constructed from gDNA extracted from fin clips for each species using a single *Sbf1* restriction enzyme as previously described^50^. Each library was sequenced on one lane of Illumina HiSeq 2500. The summary of the output of each dataset can be found in **Table S4**.

### Analysis of population genomic data for sex-specific signals

RAD-seq: Raw reads were demultiplexed with the *process_radtags.pl* script from *stacks* version 1.44^51,52^ with all parameters set to default. Demultiplexed reads for each species were analyzed with the RADSex computational workflow using the *radsex* software version 1.1.2 (10.5281/zenodo.3775206). After generating a table of markers depth with *process*, the distribution of markers between sexes was computed with *distrib* and markers significantly associated with phenotypic sex were extracted with *signif* using a minimum depth of 10 (-d 10) for both commands and all other settings to default. All figures for RAD-Seq results were generated with *sgtr*(10.5281/zenodo.3773063)^78^.

Pool-Seq: For each dataset, Pool-Seq reads were aligned to the reference genome using BWA mem (version 0.7.17)^53^. Alignment results were sorted by genomic coordinates using samtools sort (version 1.10)^54^ and PCR duplicates were removed using samtools rmdup. A file containing nucleotide counts for each genomic position was generated with the *pileup* command from PSASS (version 3.0.1b, 10.5281/zenodo.3702337). This file was used as input to compute F_ST_ between males and females, number of male- and female-specific SNPs, and male and female depth in a sliding window along the entire genome using the *analyze* command from PSASS with parameters --window-size 50000, --output-resolution: 1000, --freq-het 0.5, --range-het 0.15, --freq-hom 1, --range-hom 0.05, --min-depth 1, and --group-snps. The entire analysis was performed with a snakemake workflow available at https://github.com/SexGenomicsToolkit/PSASS-workflow. All figures for Pool-Seq results were generated with *sgtr* version 1.1.1 (10.5281/zenodo.3773063).

### Genome assembly

Raw Illumina sequencing reads for *Dallia pectoralis*, *Esox masquinongy*, *Esox niger*, and *Novumbra hubbsi* were assembled using the DISCOVAR *de novo* software^55^ with the following assembly parameters: MAX_MEM_GB=256, MEMORY_CHECK=False, and NUM_THREADS=16. Assembly metrics were calculated with the assemblathon_stats.plscript^56^ and assembly completeness was assessed with BUSCO (version 3.0.2)^57^ using the Actinopterygii gene set, and both results are given in **Table S5**.

To facilitate the comparison of sex loci in different populations of *E. lucius*, we improved the continuity of the previously published assembly of an *E. lucius* European male individual (GenBank assembly accession: GCA_007844535.1). This updated assembly was performed with Flye (version 2.6)^58^, using standard parameters and --genome-size set to 1.1g to match theoretical expectations^59^. Two rounds of polishing were performed with Racon (version 1.4.10)^60^ using default settings and the Nanopore reads aligned to the assembly with minimap2 (version 2.17)^61^ with the “map-ont” preset. Then, three additional rounds of polishing were performed with Pilon (version 1.23)^62^ using the “--fix all” setting and the Illumina reads aligned to the assembly with BWA mem (version 0.7.17)^53^. Metrics for the resulting assembly were calculated with the assemblathon_stats.pl script^56^. The assembly’s completeness was assessed with BUSCO (version 3.0.2)^57^ using the Actinopterygii gene set (4,584 genes) and the default gene model for Augustus. The resulting assembly was scaffolded with RaGOO (version 1.1)^63^ using the published female genome anchored to chromosomes (Genbank accession: GCA_011004845.1) as reference.

### Sequencing of *amha* and *amhby* genes

For species closely related to *E. lucius* (*E. aquitanicus, E. cisalpinus and E. reichertii*), the sequence of *amh* homologs was amplified and sequenced with primers designed on the sequences of *E. lucius* (**Table S6**). To search for *amh* homologs in the more divergent species (*E. niger, E. masquinongy, N. hubbsi, D pectoralis and U. pygmaea*), we generated draft genome assemblies from phenotypic males. Lastly, for *E. americanus americanus* and *E. americanus vermiculatus*, ortholog sequences were amplified with primers designed on regions that were found to be conserved in the other *Esox* species. Briefly, we first tried to amplify the complete ortholog sequences of *amha* and *amhby*, if any, from different species of Esociformes using primers designed on the exons of *amha* and *amhby* sequences of *E. lucius*. These two sequences share 78% identity, and PCR primers were chosen in divergent sequences ensuring a specific amplification of each ortholog. When the amplification was successful, the PCR products were purified using a NucleoSpin Gel and a PCR Clean-up Kit (Macherey-Nagel, Duren, Germany) and then sequenced directly by Sanger sequencing using the same PCR primers. A second round of amplification was performed using species-specific primers and primers designed on conserved regions of the *amha* and *amhby* sequences amplified in the first round. All primers used in this study were designed using Perl-Primer^64^ (Marshall 2004) and are listed in **Table S6**. The amplicons for each species were assembled with the CAP3 program^65^. Details of each approach are given in **Supplementary note 1**.

### Gene phylogeny analysis

Phylogenetic reconstructions were performed on all *amh* homolog sequences obtained from Esociformes with the *Salmo salar amh* used as an outgroup. Full-length CDS were predicted with the FGENESH+^66^ suite based on the genomic sequence and Amh protein sequence from *Esox lucius.* Sequence alignments were performed with MAFFT (version 7.450)^67^. Both maximum likelihood and Bayesian methods were used for tree construction with IQ-TREE (version 1.6.7)^68^ and Phylobayes (version 4.1)^69–71^, respectively. Details for the methods can be found in **Supplementary note 3**. The resulting phylogenetic trees were visualized with Figtree (version 1.4.1).

## Data availability

Whole Genome Shotgun genome assemblies are in the process of being deposited at DDBJ/ENA/GenBank. All gene sequences, genomic, Pool-seq and RAD-Seq reads were deposited under the common project number PRJNA634624.

## Acknowledgments

We are grateful to the fish facility of INRAE LPGP for support in experimental installation and fish maintenance and to the genotoul bioinformatics platform Toulouse Midi-Pyrenees (Bioinfo Genotoul) for providing help, computing and storage resources. This project was supported by funds from the Agence Nationale de la Recherche, the Deutsche Forschungsgemeinschaft (ANR/DFG, PhyloSex project, 2014-2016, SCHA 408/10-1, to Y.G and M.S), and the National Institutes of Health (USA) grant R01GM085318, to J.H.P). The MGX and Get-Plage core sequencing facilities were supported by France Genomique National infrastructure, funded as part of “Investissement d’avenir” program managed by the Agence Nationale pour la Recherche (contract ANR-10-INBS-09). We would like to thank MNHN curator for giving access to the collections of *Esox aquitanicus* and *Esox cisalpinus* as well as for the ONEMA and G.B. Delmastro who aided the specimen sampling. We would also like to thank Mackenzie Garvey and Penny Swanson for their assistance with dissection of *Novumbra hubbsi*.

## Author contributions

Y.G., J.H.P., M.S. obtained the funding for the project. Y.G., J.H.P., M.S., and Q.P. designed the experiments. R.F. and C.K. assembled the genomes. R.F. and Q.P. conducted analyses on RAD-Seq and Pool-Seq data. Q.P. performed genotyping and obtained all gene sequences and performed the phylogenetic analyses with the input from H.D. Q.P. and E.J. performed DNA extraction. C.R., C.E. and J.L conducted library preparation and sequencing of Nanopore long reads and Pool-Seq reads. L.J. and H.P. conducted library preparation and sequencing of RAD-Seq. Q.P., R.F., H.D., and Y.G. made the figures. J.H.P., M.S., B.K., E.R., F.W.G., W.A.L., L.B., M.T., S.S.C., E.S., G.P.J.D., F.v.H., S.C., E.A.I., V.T., H.V., J.A.L., K.O., H.H., R.G., R.T., P.D., and K.M.N. performed sample collection and determination of phenotypic sex. Q.P. wrote the manuscript with inputs from Y.G., R.F., H.D., M.S. and J.H.P. All co-authors approved the manuscript.

