## Supplemental notes, figures and tables for "The rise and fall of the ancient northern pike master sex determining gene"

### (methods, results, figures and tables)

#### Supplementary note 1. Identification of *amh* homologs in Esociformes species

To survey the presence of *amha* (the canonical copy of *amh*) and *amhby* in Esociformes, we collected samples for all species in the genus *Esox* (*E. americanus americanus, E. americanus vermiculatus, E. aquitanicus, E. cisalpinus, E. masquinongy, E. reichertii,* and *E. niger*), as well as *Novumbra hubbsi*, the only species of the genus *Novumbra*, the only well-recognized species within the genus *Dallia*, *D. pectoralis*) and one species from the genus *Umbra*, *U. pygmaea.* In total, we obtained samples from 11 species (**Table 1**). To search for *amh* homologs in the genome of these 11 species, we either 1) sequenced PCR amplicons with custom primers (see **Table S6** for primer sequences) and / or 2) sequenced and assembled the genome of some of these species (see **Table S5** for assembly metrics). For species closely related to *E. lucius* (*E. aquitanicus, E. cisalpinus,* and *E. reichertii*), *amh* homologs were amplified with primers designed specifically on *amha* or on *amhby* of *E. lucius*. Complete sequences of *amh* homologs were obtained from overlapping amplicons covering the entire genomic regions of both *amha* and *amhby* with primers anchored from upstream of the start codon (SeqAMH2Fw1 and SeqAMH1Fw1) and downstream of the stop codon (SeqAMH2Rev3 and SeqAMH1Rev4). All PCR amplicons were then Sanger sequenced from both directions and assembled to make consensus gene sequences. For more divergent species (*E. niger*, *E. masquinongy*, *N. hubbsi*, *D. pectoralis* and *U. pygmaea*), we generated draft genome assemblies (see Methods section "genome assembly", and **Table S5**) from a single heterogametic sex individual in each of these species. For *E. niger*, *E. masquinongy*, *N. hubbsi* and *D. pectoralis*, we used *amha* and *amhby* genomic sequences from *E. lucius* as queries and searched for *amh* homologs with blastn (https://blast.ncbi.nlm.nih.gov/Blast.cgi, version 2.10.0+[1]) using the parameters "Max target sequences" set to 50 and "Max matches in a query range" set to 1. This approach returned two contigs containing each of the two *amh* homologs for *E. niger* (flattened_line_76969 and flattened_line_4455), three contigs for *E. masquinongy* (flattened_line_4163, flattened_line_62015, and flattened_line_92831; while the first contig contained sequence homologous to the entire ~3 Kb of *amh,* the latter two smaller contigs contained sequences homologous to the first two Kb and third Kb of *amh* genomic sequences, respectively), two contigs in *N. hubbsi* (flattened_line_15615, and flattened_line_3346), and only one contig for *D. pectoralis* (flattened_line_2941). For *U. pygmaea*, the most divergent species from *E. lucius* in this study, blasting with *E. lucius amha* and *amhby* sequences did not yield any result. We used the same blastn strategy with the CDS of *amh* from *Salmon salar,* which returned only one contig (633485). For the other two more distantly related species of *E. lucius*, *E. americanus americanus* and *Esox americanus vermiculatus,* as no genome was available, we designed primers on regions that appeared conserved in the other *Esox* species (**Table S6**) for *amha* and *amhby* separately.

#### Supplementary note 2. Evidence supporting a single *amh* gene in *Dallia pectoralis* and *Umbra pygmaea*

We identified two *amh* homologs in all surveyed species of *Esox* and *Novumbra*. By contrast, only one *amh* homolog was found in the genomes of *D. pectoralis* and *U. pygmaea*. To provide additional evidence that this absence of a second *amh* homolog in these species was not due to assembly incompleteness, we first retrieved tissue-specific transcriptomes for both species from the PhyloFish database[2] that includes data from testicular tissue in which *amh* is usually highly expressed in teleosts[3] and in which both *amha* and *amhby* were found to be expressed in *E.*

*lucius*[4]. In agreement with our genomic data, only one *amh* transcript similar to the one identified in assemblies (100% identity over the overlapping regions) was found in the transcriptomes of each species. Besides, to verify that the two copies of *amh* were not collapsed into a single sequence during assembly, we computed the total number of heterozygous sites on the *amh* region in population genomic data (**Methods**) with the expectation that the presence of two divergent gene copies should result in high apparent heterozygosity when remapped on the single copy assembled in the genome. We compared these heterozygosity data in *D. pectoralis* and *U. pygmaea* to similar results observed when mapping sex-specific pooled libraries of *E. lucius* (N=30 males and 30 females, respectively) to a female reference genome containing only *amha* (GCA_004634155.1). If there is only one copy of the *amh* gene in *D. pectoralis*, we expect a low level of heterozygosity on the *amh* genomic region, comparable to what is obtained after mapping female Pool-Seq reads of *E. lucius* to a reference genome with only one *amh* gene. On the other hand, if two divergent *amh* genes exist in *D. pectoralis* while only one is present in the reference genome, then the heterozygosity of the *amh* genomic region would be similar to what is obtained after mapping male *E. lucius* Pool-Seq reads to a female reference genome. Reads from the male and female pools of *E. lucius* and from male pool of *D. pectoralis* were aligned separately to the reference genome (GCA_004634155.1) and our draft genome, respectively, using BWA mem version 0.7.17[5] with default parameters. Each resulting BAM file was sorted and PCR duplicates were removed using Picard tools version 2.18.2 (http://broadinstitute.github.io/picard) with default parameters. Afterwards, variants were called for each BAM file using bcftools mpileup version 1.9[6] with default parameters on genomic regions containing *amh*, which locates between 12,906,561 bp and 12,909,640 bp on LG08 of *E. lucius* (GCA_004634155.1: CM015581.1), and between 23,447 bp and 25,910 bp on the flattened_line_2941 contig of the draft genome of *D. pectoralis*. In total, 106 variants were observed in the pool of *E. lucius* males on the *amha* region, resulting from the mapping of reads originating from both *amha* and *amhby,* while only 12 variants (true allelic variations) were observed in the same region when mapping reads from the female pool originating only from *amha*. With male Pool-Seq reads from *D. pectoralis*, we observed only four variant sites on the ~3 Kb *amh* region, supporting that only one *amh* gene is present in the genome of *D. pectoralis*. Although sex-specific Pool-Seq reads were not available for *U. pygmaea*, we performed the same analysis with reads from the single male individual used to assemble the genome. No variant was observed in the ~3 Kb region containing *amh* located between 760,763 bp and 757,962 bp on contig 633485 of our draft genome of *U. pygmaea*, supporting that only one *amh* gene is present in this species.

#### Supplementary note 3. Phylogenetic relationships among amh homologs.

To clarify the relationship among *amh* homologs and estimate the duplication time of *amh* in Esociformes, different phylogenetic analyses were carried out in addition to the Maximum likelihood method (ML) analysis with all coding sequences (CDS) sequences presented on **Figure 2A**. Additional analyses were carried with or without the truncated *amh* / Amh sequences of *E. niger* and *N. hubbsi* (see section 6) using the *amh* / Amh homolog of *Salmo salar* as an outgroup. CDS and proteins were predicted with the FGENESH+ [7]suite, based on their genomic sequences. These putative CDS and protein sequences were then aligned using MAFFT (version 7.450)[8], residue-wise confidence scores were computed with GUIDANCE 2[9] and only well-aligned residues with confidence scores above 0.99 were retained. The resulting alignment file was used for model selection and tree inference with IQ-TREE (version 1.6.7)[10] with 1000 bootstraps and 1000 SH-like approximate likelihood ratio test for robustness. In addition, to verify that the topology of the single *amh* homolog of *D. pectoralis* was not an artefact of long branch attraction, we also constructed phylogenetic trees with only the first and second codons of the coding sequences[11] as well as with complete protein sequences. For an additional confirmation for the topology of trees we obtained for the *amh* homologs, the same alignments were run in a Bayesian framework implemented in Phylobayes (version 4.1)[12–14] using the CAT-GTR model with default parameters, and two chains were run in parallel for approximately 1000 cycles until the average standard deviation of split frequencies remained ≤ 0.01.

In all analyses, two divergent monophyletic clusters containing the same sequences were consistently observed (**Figure S1**) in agreement with the results obtained with the ML methods on all CDS (**Figure 1A**). The first cluster contained the *amha* sequence from *E. lucius* and one sequence from all other surveyed species of Esocidae (*Esox*, *Dallia* and *Novumbra*), while the second cluster contained all the remaining *amh* homologs from species with two *amh* sequences including the *amhby* sequences of *E. lucius* (**Figure 1A and Figure S1**). The topologies within each group were in agreement with the Esociformes species tree (**Figure 1B**), suggesting that the duplication of *amh* occured in a common ancestor of the Esocidae and has been maintained through speciation events[15]. Based on these results, we identified sequences in the two clusters as *amha* and *amhby* orthologs, respectively. The single *amh* ortholog of *U. pygmaea,* the closest sister species to *Esocidae*, fell outside of the *amha* / *amhby* clade. By contrast, the single *amh* ortholog of *D. pectoralis* was consistently placed within the *amha* cluster with a strong bootstrap support (96.8% / 98%) whatever the approach chosen (**Figure S1**), and was genetically more similar to other *amha* sequences than to *amhby* duplicates (77. 8% and 80.9%) indicating that this sequence is orthologous to *amha.* These results strongly suggest that the placement of the single *amh* ortholog of *D. pectoralis* within the *amha* cluster is not an artefact of tree reconstruction, that the duplication of *amh* most likely occurred after the split of the *Umbridae* and *Esocidae* families between 65 and 88 Mya[16] and that *amhby* was subsequently lost in *D. pectoralis*.

#### Supplementary note 4. Association between *amhby* and male phenotype

To determine if male sex-linkage is a general feature of *amhby* in Esociformes, we tested the association between the presence of *amhby* and male phenotype in samples with known phenotypic sex. A significant sex-linkage with *amhby* present in 100% of the male samples and 0% of the female samples was found in *E. americanus*, *E. reichertii*, and *E. niger* (**Table 1**), indicating that sex is determined by an XX/XY system with *amhby* in the sex locus for these three species. In *E. cisalpinus* and *E. aquitanicus*, *amhby* was also found in all male samples but in none of the female samples, suggesting that *amhby* is also male specific in these species, but low sample sizes (two individuals of each sex) prevented statistical testing. Overall, these results show that *amhby* may be a conserved master sex-determining gene in most Esocidae (*E. lucius*, *E. americanus, E. reichertii, E. niger, E. cisalpinus,* and *E. aquitanicus*). In *E. masquinongy*, *amhby* was significantly associated with male phenotype in two populations (Iowa and Wisconsin, US). However, in a third population from Quebec, Canada, the association was not significant (**Table 1**). Genotypes were also not compatible with the expected hemizygous status of a Y sex locus as *amhby* was found heterozygous in two males and one female (**Figure 1**). Variation in sex determination in populations of *E. masquinongy* and *E. lucius* is analyzed in more detail below in Supplementary note 8. In *N. hubbsi*, *amhby* was found in 21 out of 23 males (95%) and only one out of 22 females (4%), showing significant association with male phenotype which is also indicative of an XX/XY SD system.

#### Supplementary note 5. No signature of relaxation from purifying selection of *amha* and *amhby* genes

To compare selection pressure between *amha* and *amhby* genes, dN/dS ratios were computed between each ortholog sequence and the *amh* sequence of *U. pygmaea*, which was found to have diverged from the other Esociformes prior to the *amh* duplication. Sequences from *E. niger* and *N. hubbsi* were excluded as their *amhby* orthologs were substantially shorter and could introduce bias in the analysis. The ratio of nonsynonymous to synonymous substitution (dN/dS = $\omega$) among *amha* and *amhby* sequences was estimated using PALM (version 4.7)[17] based on aligned full length CDS, and the phylogenetic tree obtained with the CDS was used in the analysis. First, a relative rate test on amino-acid substitution[18] was performed between *amha* and *amhby* pairs with the *amh* of *U. pygmaea* used as an outgroup sequence: for each species with *amh* duplication, $\omega$ was calculated between the *amha* ortholog and *amh* of *U. pygmaea*, and was compared to the $\omega$ calculated between the *amhby* ortholog and *amh* of *U. pygmaea*. A Wilcoxon test was used to compare the resulting $\omega$ values for the *amh* paralogs of these species. Then, several branch and site models (M0, M1a, M2a, M7, M8, free-ratio, and branch-site) implemented in the CODEML package were fitted to the obtained data. The goodness of fit of these models was compared using the likelihood ratio test implemented in PALM. The resulting dN/dS ratios (**Table S7**) were not significantly different between the two *amh* paralogs, with a mean of 0.378 for *amha* and a mean of 0.376 for *amhby* (Wilcoxon signed-rank test p=0.9015). In order to identify potential positive selection signals on specific sites, several site models allowing different dN/dS ratios among sites were used. However, none of these alternative models fitted the data better than the null model, which allowed a single dN/dS for all sites and all branches in the gene phylogenetic tree of *amh* orthologs **(Table S8**). In summary, we did not detect signature of either post-duplication relaxation of purifying selection or directional selection on either *amha* or *amhby* orthologs without structural variation.

#### Supplementary note 6. Truncations of Amhby in *E. niger* and *N. hubbsi*

The typical *amh* gene in teleosts comprises seven exons encoding for a protein that contains 500 to 571 amino acids with a C-terminal TGF-β domain that is crucial for canonical Amh function[3,19]. The predicted structures of the Esociformes Amh homologs generally agree with this classical teleost Amh structure. In *E. niger* and *N. hubbsi*, however, Amhby proteins appear to be truncated. In *E. niger*, the conserved 5' and 3' terminal exons were missing from the genome assembly and the predicted *A*mhby protein would contain only three of the conserved Amh exons (exon5, exon6 and a partial exon7 with a truncated TGF-β domain). In addition, the *E. niger amhby* genomic sequence is flanked by repeated elements (55 blastn results with e-value < 9e-16 and 250 blastn results with e-value < 2e-10 for the element upstream and downstream of *amhby*, respectively) suggesting that this truncation was caused by repeat insertions, a phenomena predicted and observed to occur on sex chromosomes due to reduced efficacy of selection[20–25]. In *N. hubbsi*, the predicted Amhby protein contains six of the seven conserved exons found in other Esocidae Amhby proteins with a predicted first exon encoded by only 8 amino acids sharing no homology with the 50 amino acid sequences of the first Amhby exon of other Esocidae. This Amhby first exon is conserved among all non-truncated Esocidae Amhby sequences but shares no homology with other teleost Amh sequences (**Figure S3A**). In *N. hubbsi*, this small predicted exon1 was found 4 Kb upstream of the second exon and no homologous sequence to the first conserved Amh exon of *Esocidae*, which typically lies about 1 Kb upstream of the second exon, was found in this region. Supporting this result, a similar gene model was predicted from another *N. hubbsi* male genome assembly, sequenced with long-reads and with much better genome continuity (Simen Rod Sandve, Michel Moser and Sigbjorn Lien, personal communications). To investigate whether these truncations could be assembly artifacts, we searched for potential missing homologous sequences in raw genome reads of both species using tblastn (https://blast.ncbi.nlm.nih.gov/Blast.cgi, version 2.10.0+[1]) using the exon1 protein sequence from *E. lucius* as a query. For both species, no homologous sequence (with an e-value < 1E-10 ), besides the corresponding exons from *amha*, was obtained, indicating that truncation of these segments of *amhby* in both species is not an artefact of assembly.

#### Supplementary note 7. Sex determination systems in *Novumbra, Dallia and Umbra*

Apart from *E. lucius*, SD systems were not described for most Esociformes. As our *amhby* sex-linkage results suggest that XX/XY SD system is common among *Esox* species. To complement previous results in *E. lucius*[4] and sex-linkage analyses we performed Restriction Associated DNA Sequencing (RADs-Seq) (see Methods) in *N. hubbsi* (24 females, 26 males), and *U.*

*pygmaea* (31 individuals for each sexes) as well as RAD-Seq and Pool-Seq in *D. pectoralis* (30 individuals for each sexes).

In *N. hubbsi*, *amhby* was found to be significantly associated with male phenotype, indicating an XX/XY sex determination system. Two of the 23 males did not carry *amhby*. It is worth noting that the phenotypic sex of the samples used in this study was solely determined based on the external phenotype, as *N. hubbsi* is endemic to western Washington state (USA) and is considered a state sensitive species[26]. As a result, these two male outliers could be individuals with misassigned phenotypic sex. To complement the *amhby* association results, we generated RAD-Seq reads for 26 males and 24 females *N. hubbsi* collected from Olympic National Park (Washington, USA). We identified two markers that were present in 19 (80%) males and one female, however, their association with sex phenotype was not significant after Bonferroni correction. Although all our samples were from the same large region (4000 $km^2$), they were collected from six different locations (James Pond, Powell Creek, Stearns Wetland, Conner Creek, Steamboat Bog and S. Hanaford Creek). Previous studies revealed very limited gene flow among these locations and that different collection locations represented genetically distinct populations[26]. We therefore hypothesized a strong population structure and potential sex locus polymorphism among different populations could weaken the sex-specific signal.

To test for population structure in our dataset, we estimated $F_{ST}$ for each pair of locations using STACKS[27,28]. RAD-Seq reads were demultiplexed with the *process_radtags.pl* wrapper script and analyzed with *denovo_map.pl* using the following parameter values: m=10, M=2, N=3, "--gapped" not set, and "H" not set. Pairwise $F_{ST}$ was computed from the results of *denovo_map.pl* with the *populations* pipeline with minimum percentage of individuals in a population set to 0.5 (r=0.5), minimum stacks depth at a locus set to 10 (m=10), and minimum populations at a locus set to 2 (p=2) and the resulted $F_{ST}$ values are presented in **Table S9**. The S. Hanaford Creek and Stearns Wetland populations had the lowest pairwise $F_{ST}$ (0.0031) and were highly divergent from all other populations ($F_{ST}$ ranging from 0.21 to 0.58). Coincidentally, the two markers associated with male phenotype that we identified in our RADSex analysis on all samples were absent in males from these two populations. Among the other four populations, these two markers (out of 53126) were significantly associated with male phenotype (p=0.021, chi- squared test with Bonferroni correction) and no marker was significantly associated with female phenotype, agreeing with the XX/XY SD system from the results of *amhby* association. One hypothesis for the absence of these male-specific markers in the Hanaford Creek and Stearns Wetland populations is that in these two populations the Y locus could be smaller or contain different sequences without *SbfI* cleavage sites. Overall, our results indicate an XX/XY SD system in *Novumbra hubbsi* with a small sex locus about 48 Kb (**Supplementary note 10**) containing *amhby*, and that at least two Y haplotypes differing in size and/or sequence are present among the surveyed populations.

In *D. pectoralis*, 48,475 markers were identified in the RADSex analysis, and among them, one marker was found in 25 of 30 females (83%) but in only three males (10%), and another marker was found in 27 females (90%) and 8 males (27%). After applying Bonferroni correction, one marker was significantly associated with female sex (Chi-squared test, p = 0.0027) and no marker was significantly associated with male sex (**Figure 3B**). These results suggest a ZZ/ZW SD system in *D. pectoralis* with a small sex locus around 65 Kb (**Supplementary note 10**). To investigate the SD system and potentially the sex locus with higher resolution in this species, we generated Pool-Seq reads using a male and a female pool comprised of the same individuals. Because the genome we produced for this species was too fragmented for a reference-based analysis, we adopted a reference-free approach searching for sex-biased k-mers in the Pool-Seq data. First, 31-mers were identified and counted in the male and female pools reads using the "count" command from Jellyfish (version 2.2.10[29]) with the option "-C" activated to count only canonical k-mers and retaining only k-mers with an occurence higher than five and lower than 50,000. Afterwards, tables of k-mer counts produced by Jellyfish were merged with the "merge" command from Kpool (https://github.com/INRA-LPGP/kpool) and the resulting merged table was filtered using the "filter" command to only retain kmers present more than 25 times in one sex and less than 5 times in the other sex; such k-mers were considered sex-biased. Using this criterion, we identified 45 times more female-biased k-mers in (N = 1,081,792) than male-biased k-mers (N = 23,816), indicating that females likely carry genomic regions that are absent from the males and thus are the heterogametic sex[30]. The clear overabundance of female-biased k-mers confirms the results of our RADSex analysis indicating that a ZZ/ZW sex determinations system is operating in *D. pectoralis*.

The Umbridae diverged from the rest of the Esociformes before the duplication of *amh* and the recruiting of *amhby* as MSD gene, and therefore species from this clade are likely to have a different MSD gene. To date, nothing is known about sex determination in this clade, and characterizing the sex-determination mechanism of the *Umbridae* is a crucial step towards determining the potential ancestral MSD of this clade. To initiate this process, we generated RAD-Seq reads for 31 males and 31 females from one of the three *Umbra* species, *Umbra pygmaea*. Using the RADSex workflow, we identified 140 markers significantly associated with male sex and no markers associated with female sex out of 6,501,900 total markers. These results clearly indicate an XX/XY sex determination system with a rather large non-recombining region around 5.4 Mb (**Figure 3C, Supplementary note 10**).

#### Supplementary note 8. Transitions of sex determination systems among different populations of *E. masquinongy*

In *E. masquinongy*, sex-linkage genotyping revealed different frequencies of males and females carrying *amhby* in different populations **(Table 1)**. Beside the absence of *amhby* sex-linkage in the Quebec population, we also identified two males and one female (among 20 males and 20 females) being apparent heterozygous for *amhby* with a clear bi-allelic SNPs in sanger sequencing results (**Figure S2**). To investigate whether these differences originate from variation in SD systems, we generated RAD-Seq data for 25 males and 28 females from the Iowa (US) population, where *amhby* was significantly associated with male phenotype (Chi-squared test, p-value = 2.01e-8), and for 20 males and 20 females from the Quebec (Canada) population, where the association was not significant (Chi-squared test, p-value = 0.18). These datasets were analyzed with the RADsex workflow[31] with a minimal depth of 5 (see Materials and Methods). In the Iowa population, 5 markers among a total of 308,960 were significantly associated with males while no marker was associated with females (**Figure 3E**). This result indicates an XX/XY sex determination system with a small non-recombining sex locus, around 130 Kb (**Supplementary note 10**), as observed in *E. lucius*[4] and *N. hubbsi* (see **Supplementary note 7**). In contrast, no sex-specific marker was identified in the Canadian population lacking *amhby* association (248,698 total markers, **Figure 3D)**.

To better characterize the sex locus of *E. masquinongy*, we generated reads from a male pool and a female pool using individuals from the Iowa population for which RADSex found an XX/XY SD system. We sequenced and assembled the genome of a male individual from this Iowa population using Illumina short reads (see Materials and Methods) and scaffolded the draft assembly into chromosomes based on homology with the North American *E. lucius* reference genome (GCA_004634155.1) using RaGOO version 1.1[32]. We then used the PSASS workflow (https://github.com/SexGenomicsToolkit/PSASS-workflow) to align reads from each pool to the assembly and to compute 1) the number of male-specific SNPs in 50 Kb non-overlapping windows, and 2) the depth of aligned reads for each sex in 1 Kb non-overlapping windows to identify regions showing differentiation between sexes, setting both "window-size" and "output-resolution" to 50 Kb or to 1 Kb and "group-snps" to 'True'. First, we found that the 3 Kb region containing the genomic sequence of *amhby* (unplaced_scaffold_RaGOO_chr0: 24,967,172- 24,968,938) is covered by only male reads, confirming the tight association between the *amhby* ortholog and male phenotype in *E. masquinongy* (**Figure S5.A**). Second, we found the highest number of male-specific SNPs (MSS) in a single window located around 100-150 Kb on LG24 that contained 212 MSS and zero female-specific SNPs (**Figure S5.B, Figure S5.C**), while genome average was 1.57 MSS per 50 Kb window (3.17 MSS per window when excluding 50 Kb windows without MSS). This result indicates that the sex locus in *E. masquinongy* is homologous to the proximal end of LG24 in *E. lucius*. Besides LG24, no other linkage group contained a window enriched with MSS (**Figure S5.B**). Overall, this Pool-Seq analysis confirms the XX/XY SD system with a low differentiation between the X and Y chromosomes identified with the sex linkage and RAD-Seq analyses, as observed in *E. lucius*[4] and *N. hubbsi* (see above), and supports the hypothesis that the sex locus of *E. masquinongy* is similar to that of *E. lucius*.

#### Supplementary note 9. The recent loss of *amhby* in North American populations of *E. lucius* is associated with the absence of a well-differentiated genetic sex determination system

Although *amhby* was demonstrated functionally as the MSD gene in European populations of *E. lucius* (Pan et al., 2019), the *amhby* sequence is absent from the genome assembly of a male specimen from a Canadian population (GCA_000721915.3)[33]. To explore whether there is population variation in sex determination, we obtained samples with known phenotypic sex from mainland USA (1 population) and Canada (3 populations). In all these North-American populations, *amhby* could not be amplified in males (N= 88) or in females (N=74), neither with primers designed specifically on *E. lucius amhby* nor with primers anchored on regions with high sequence conservation among several Esociformes species. This result suggests a complete loss of this ancestral MSD gene in *E. lucius* populations from mainland USA and Canada. To delineate the geographic distribution of northern pike populations that lost *amnby*, we further assessed its presence in Alaska, USA (two populations) and Xinjiang, China (one population), two localities that connect the European populations among which *amhby* is present and the North American populations among which *amhby* is absent. Significant association between *amhby* and sex phenotypes were confirmed in samples from these two new populations (**Table 1**), indicating that the loss of *amhby* was likely restricted to populations found in mainland USA and Canada.

To investigate whether the loss of *amhby* coincides with large genomic changes in Canadian and USA *E. lucius* populations, we generated Pool-Seq reads from a pool of males and a pool of females from a Canadian population without *amhby* (Quebec) and from a European population carrying *amhby* (Ille-et-Vilaine, France), using 30 males and 30 females for each species. In order to facilitate reference-based Pool-Seq analyses, in particular to potentially increase the continuity of the sex locus in the assembly, we generated additional Nanopore reads from the European male *E. lucius* sequenced in ou previous study[4] and re-assembled the genome. The new assembly had greatly improved continuity (scaffold N50 of 10.7 Mb compared to 800 Kb in the previous assembly) and improved complete BUSCO score (94.1% against 93.2%) (**Table S1**). Furthermore, after scaffolding with Ragoo, all Y-specific and Y-differentiated regions were located on Chromosome 24 (NCBI accession number SDAW00000000). Pool-seq reads from the Canadian and European populations were aligned on this new assembly to explore the extent of differences between male-specific regions. We computed the number of male-specific SNPs in 50 Kb non-overlapping windows and the absolute depth of mapped reads as well as relative depth to genome average of mapped reads per sex in 1 Kb non-overlapping windows across LG24 to identify regions showing differentiation between sexes with the PSASS workflow (https://github.com/SexGenomicsToolkit/PSASS-workflow) with either output window-size and output-resolution set to 50 Kb or 1 Kb, and group-snps set to 'True'.

In the European population containing *amhby*, there is a genome average of 1.7 male-specific SNPs per 50 Kb window (3.9 male-specific SNPs per window when excluding windows containing zero male-specific SNPs). In contrast, we found a single 50 Kb window containing 182 male-specific SNPs at the proximal end of LG24 (from 0.95 Mb to 1.0 Mb), showing the greatest differentiation between males and females in the genome. We then searched Y-specific regions using reads depth in 1 Kb window with few or no female reads mapped (< 3 reads per Kb) and male reads mapped at a depth close to half of the genome average (relative depth between 0.4 and 0.6) value. Based on these depth filters we identified on LG24, 70 potential Y-specific 1 Kb windows located between 0.76 Mb to 0.86 Mb and 70 Y-specific 1 Kb windows located between 1.0 Mb and 1.24 Mb, planking the male-specific SNPs enriched regions (**Figure 4A.1**). These results are consistent with previous finding using a female genome assembly as reference for mapping[4] regarding the location of the sex locus at the proximal end of LG24. However, the improved continuity of the male assembly allowed us to anchor the male-specific sequences, which were unplaced in the previous analysis, on the Y chromosome and show that the region with the highest X/Y differentiation is flanked on both sides by Y-specific sequences.

In contrast, in the Canadian population we did not found any male-specific SNPs enrichment at the proximal end of LG24 where the sex locus is located in European populations (**Figure 4A.2**). Furthermore, virtually no males or females reads were mapped to the 140 Kb Y-specific regions identified in the European population (**Figure 4B.2**) and no sign of differentiation between males and females could be observed along the rest of LG24. Altogether, these results suggest that the Canadian population and likely the mainland USA populations lack the entire sex locus found in European populations, including the MSD gene *amhby*.

To investigate whether these is a new GSD system in North American *E. lucius* populations, we searched for loci associated with sex in two independent populations from Quebec (Canada) using a combination of RAD-Seq, that is a robust approach to analyze individual variations and Pool-Seq, which increases the resolution of analyses to the detection of small genomic regions. Sex-linkage was first explored in each population using RAD-Seq data in a reference-free approach using the RADSex pipeline with default parameters and the minimal read depth set to 5**.** In both populations, we did not find any sex-linked markers among a total of 8,440,899 and 4,526,552 markers identified in each population (**Figure 5A & 5B**), indicating that if there is a new sex locus in these populations its detection escapes the resolution of RAD-Seq (31.8 RAD markers per Mb on average, see **Supplementary note 10**) due to a very low differentiation between the new sex chromosome pair. Next, we analyzed sex-specific Pool-Seq data for the Canadian population to search for regions enriched of male- or female-specific SNPs and/or regions with sex-specific read depth. Pool-Seq was analyzed with the PSASS pipeline with the same parameters as the previous Pool-Seq analysis of this dataset, but reads were mapped in this analysis onto the reference genome of a female *E. lucius* from North American (GenBank assembly accession: GCA_004634155.1). We searched for 50 Kb non-overlapping windows enriched with either male or female specific SNP. Overall, the level of differentiation between males and females was low, and the highest Fst observed across the entire genome is 0.07 located between 0.397 Mb and 0.398 Mb on linkage group 11. Very low amount of sex-specific SNPs were found for both male and female pools in this population, especially when comparing to the same analysis performed with data from an European population (**Figure S7**). In the European population, we found a genome average of 3.5 male-specific and 2.7 female-specific SNPs per 50 Kb window, with peak windows containing 625 male-specific SNPs near the LG24 sex locus, and 217 female-specific SNPs at the proximal end of LG5. In comparison, the Canadian pools had roughly similar values of average genome sex-specific SNPs (3.3 male-specific and 2.9 female-specific SNPs per window), but with peak windows of sex-specific SNPs of 49 and 45 for males and females, respectively. Given this low amount of differentiation between the sexes, and that no particular chromosome was enriched with windows containing a high amount of sex-specific SNPs, no region stood out to be the candidate region for the sex locus. Besides that, no 1 Kb window with sex-specific read depth was found. Overall, no clear signal of a sex locus was observed across the entire genome supporting the hypothesis that the mainland USA and Canadian populations lack a well-differentiated sex determining region.

#### Supplementary note 10. Resolution of RAD-Seq in Esociformes species

To help estimate the size of the sex locus from sex-specific RAD markers, we determined the number of potential *SbfI* cleavage sites based on our draft genome assemblies for each species. For *E. lucius* (Canadian population), *E. masquinongy* (Iowa population), *N. hubbsi* and *D. pectoralis*, we predicted the number of RAD-Seq cleavage sites present in each genome by counting the number of unambiguous matches for sequence of *SbfI* (CCTGCAGG), the restriction enzyme used in RAD-Seq library preparation[34]. On average, we found 31.8 RAD markers per Mb in *E. lucius* , 38.4 in *E. masquinongy* , 41.6 in *N. hubbsi*, 30.8 in *D. pectoralis,* and 26.2 in *U. pygmaea* (**Table S10**). Because we do not see large species differences (26 to 40 RAD markers / Mb) this suggests that, apart from potential local variations of the RAD markers density in sex loci, our RAD-Seq comparative analysis could to some extent be used to compare sex locus size within species on the basis of this number of sex-specific RAD markers. We are aware of the limitation that using the number of sex-specific marker usually lead to an overestimation of the size of the sex locus. For all of our species species with the exception of *U. pygmaea*, we identified very few sex-specific markers, indicating very small sex locus. This simple calculation is only intended to helps provide an rough approximation of the size of the sex locus.

### Supplementary figures

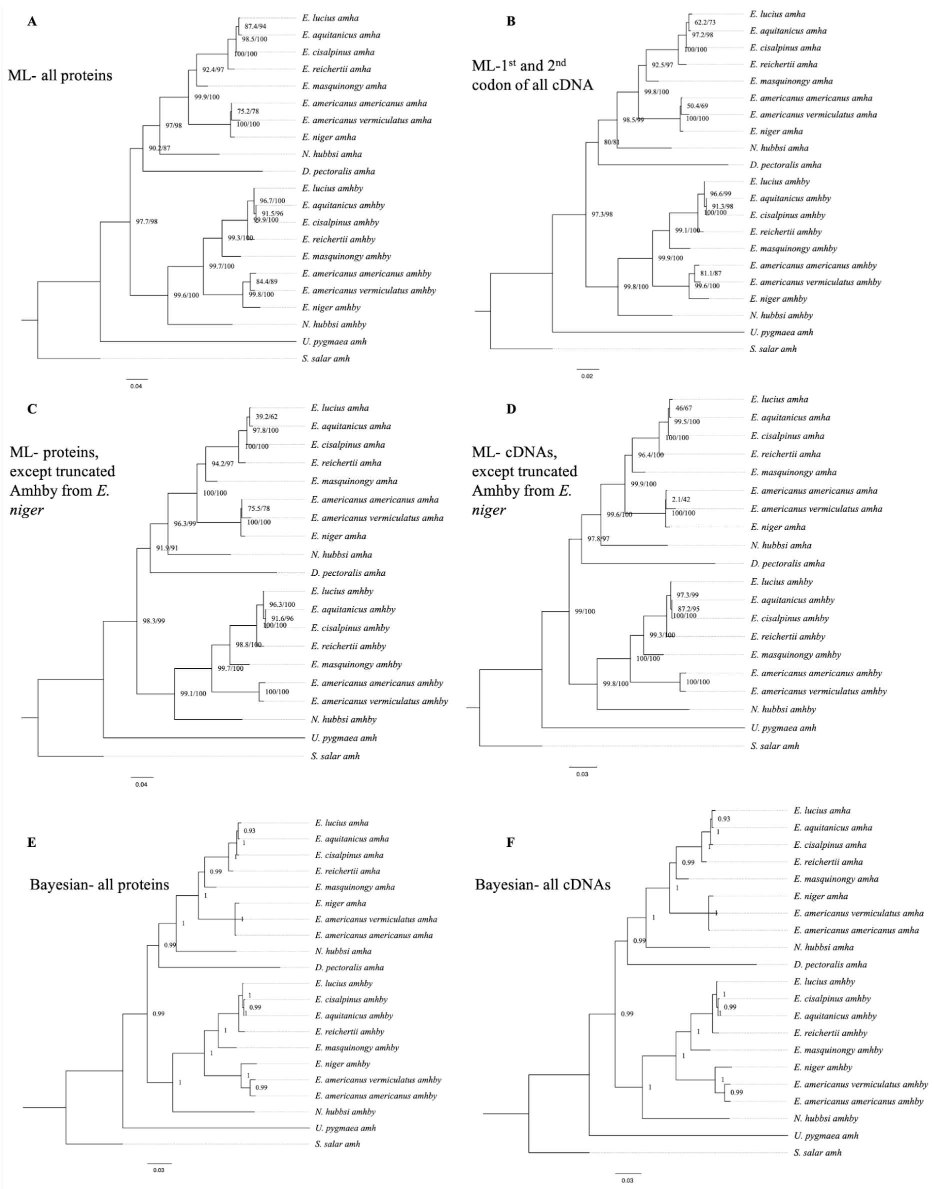


**Figure S1: Additional phylogenetic reconstruction of *amha* and *amhby* orthologs from the *Esociformes* with amh sequence from *Salmo salar* as an outgroup. A**: Phylogenetic tree built by Maximum likelihood method implemented in IQ-TREE putative protein sequences of all identified *amh* homologs. **B:** Phylogenetic tree built by Maximum likelihood method with only the 1st and 2nd codon of putative coding sequence of all identified *amh* homologs. **C**: Phylogenetic tree built by Maximum likelihood with putative protein sequences of *amh* homologs without one highly truncated sequence from *E. niger*. **D**: Phylogenetic trees built by Maximum likelihood with putative coding sequences of *amh* homologs without one highly truncated sequence from *E. niger.* **E**: Phylogenetic tree built by Bayesian method implemented in PhyloBayes with putative protein sequences of all identified *amh* homologs. **F**: Phylogenetic tree built by Bayesian method with putative coding sequences of all identified *amh* homologs. Bootstrap values are given on each nod of the tree and all trees are rooted with *amh* sequence from *S.salar*.

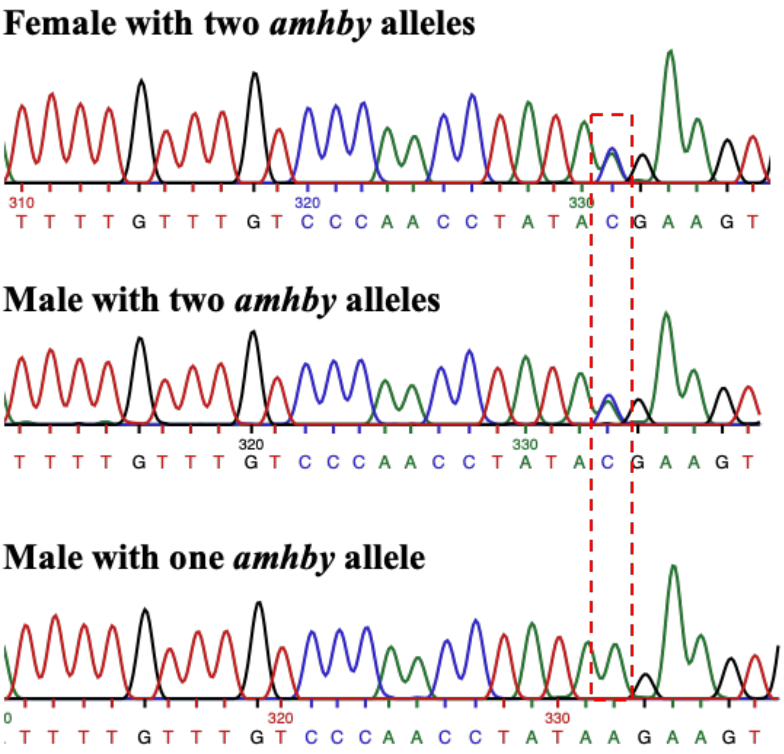


**Figure S2**. **Sanger sequencing results for *E. masquinongy* one female and one male carrying two different alleles of *amhby* and one male carrying only one *amhby* allele.** The base with a bi-allelic SNP is highlighted with the red dashed-line box.

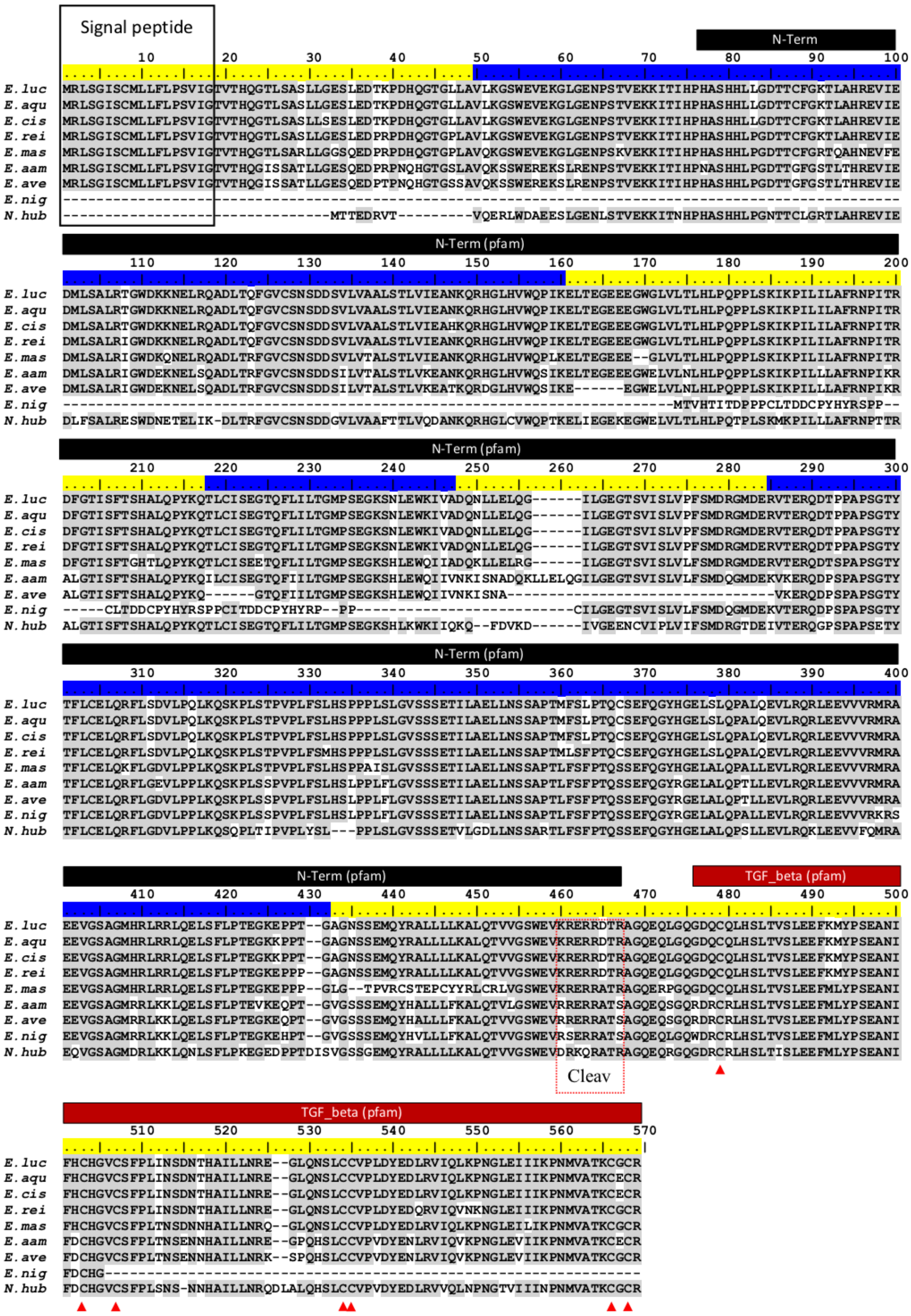

**Figure S3**. ClustalW[35] alignment of Esocidae Amhby putative protein sequences. The signal peptide was predicted with SignalP[36]. No signal peptide was detected in the Amhby sequences of both *E. niger* and *N. hubbsi*. N terminal region (indicated by a black bar) and the Transforming growth factor beta like domain (indicated by a red bar) were predicted with the Motif Scan tool at ExPASy[37] with the Pfam motif database[38] optimized for local alignments (pfam-fs, Pfam 32.0, September 2018). The seven exons of *amhby* are shown by the alternating blue and yellow colors on the sequence ruler. The seven conserved cysteines of the TGF-beta domain are indicated by red arrowheads. The region containing the putative Amh cleavage site (Cleav) is boxed in red. *E. luc* (*Esox lucius*), *E. aqu* (*E. aquitanicus*), *E. cis* (*E. cisalpinus*), *E. rei* (*E. reichertii*), *E. mas* (*E. masquinongy*), *E. aam* (*E. americanus americanus*), *E. ave* (*E. americanus vermiculatus*), *E. nig* (*E. niger*) and *N. hub* (*Novumbra hubbsi*).

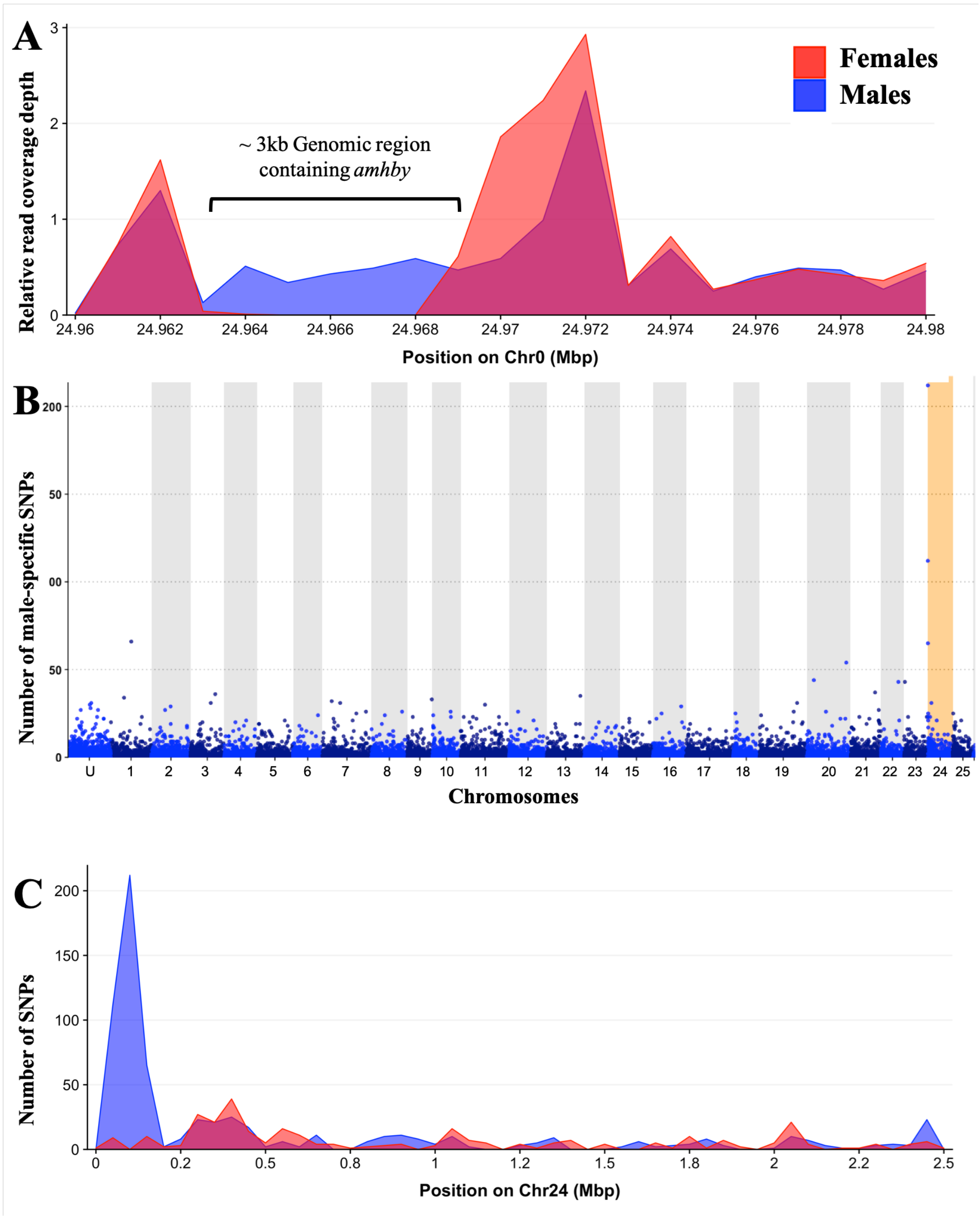


**Figure S4**: **Analysis of sex differentiation in the genome of a male *E. masquinongy* (*Ascension number*) with Pool-Seq data**. **A**: The relative to genome average coverage depth of male and female Pool-Seq reads on the region containing *amhby.* The male data are represented in blue and female in red. The 3kb region containing *amhby* is covered by virtually no female reads and by male reads at a depth about half of the genome average depth. **B**: Number of male-specific SNPs in 50kb non-overlapping windows in each linkage group in the male genome of *E. masquinongy*. The window containing the highest number of male-specific SNPs on LG24 is highlighted. **C**: Number of male and female-specific SNPs from Pool-Seq in 50 kb non-overlapping windows is plotted along LG. The window located between 0.1 to 0.15 Mb showed the strongest differentiation between males and females.

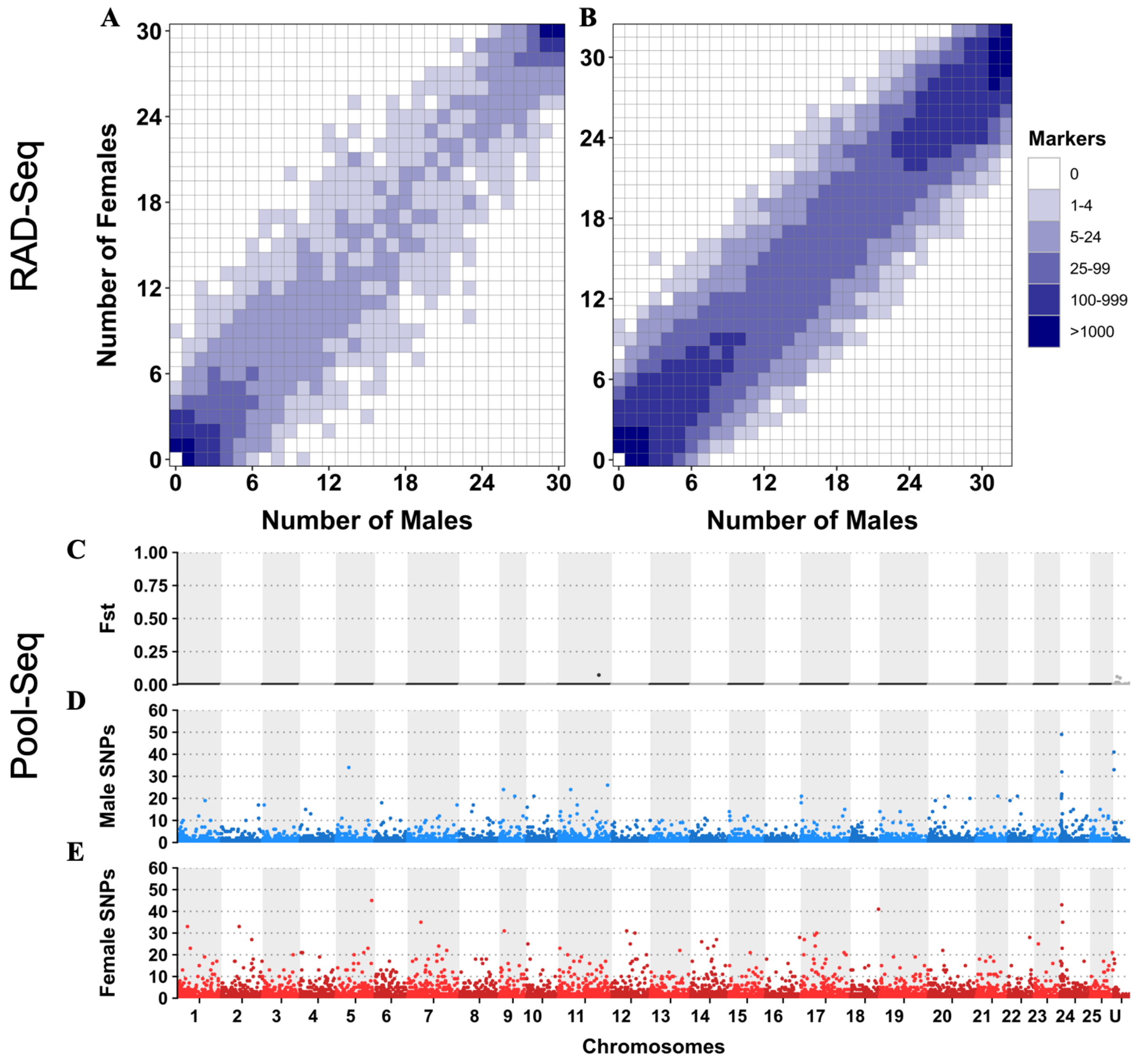


**Figure S5: No differentiation between male and female genomes revealed by RAD-Seq and Pool-Seq in some North American populations of *E. lucius*. A-B**: Distribution of RADSex markers in males and females of two populations of *E. lucius* from Canada. The distribution of markers in males and females was computed with RADSex with a minimum depth of 5 to consider a marker as present in an individual for both dataset. In each tile plot, the number of males and number of females are represented on the horizontal and vertical axes respectively, and intensity of color of a tile indicates the number of markers present in the corresponding number of males and females. We did not find any marker significantly associated with male or female sex in these two populations. **C-E**: Analysis of male/female differentiation across Canadian *E. lucius* genome with Pool-Seq reads from 30 males and 30 females mapped to the reference genome (Accession number: GCA_004634155.1). Between sex $F_{ST}$ (**C**), female specific SNPs (**D**) and male-specific SNPs (**E**) are computed for 50Kb non-overlapping windows across the 25 linkage groups (LGs) and unplaced scaffolds. The window with the highest between-sex $F_{ST}$ value of 0.07 is located on LG11 and no one particular chromosome is enriched with windows contained high numbers of male- or female- specific SNPs.

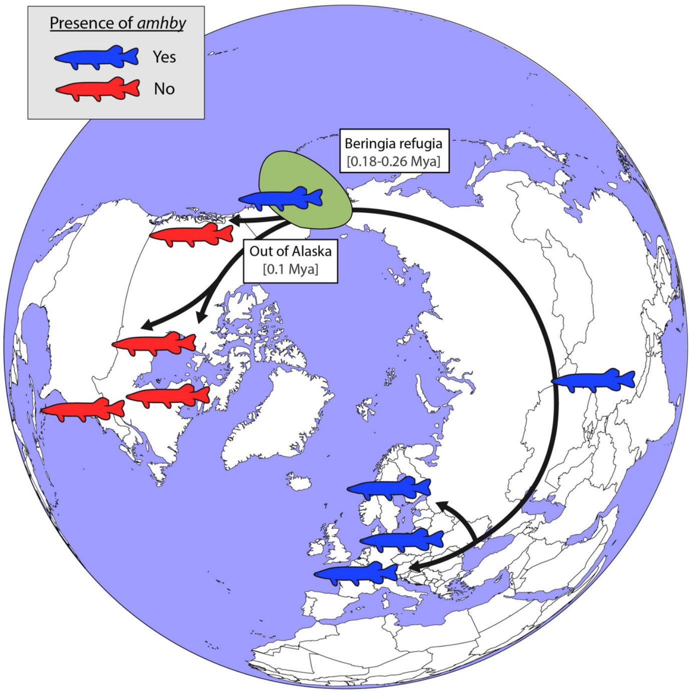


**Figure S6**: Schematic representation of the alternative hypothesized route of post-glacial *E. lucius* expansion from an Eurasian refugia ~ 0.18 to 0.26 Mya and the presence (red silhouettes) / absence (blue silhouettes) of *amhby* in different populations, showing that this master sex determining gene was lost in North American population during this out of Alaska dispersal around 0.1 Mya. The hypothesized refugia in the Beringia region is indicated with a yellow highlight.

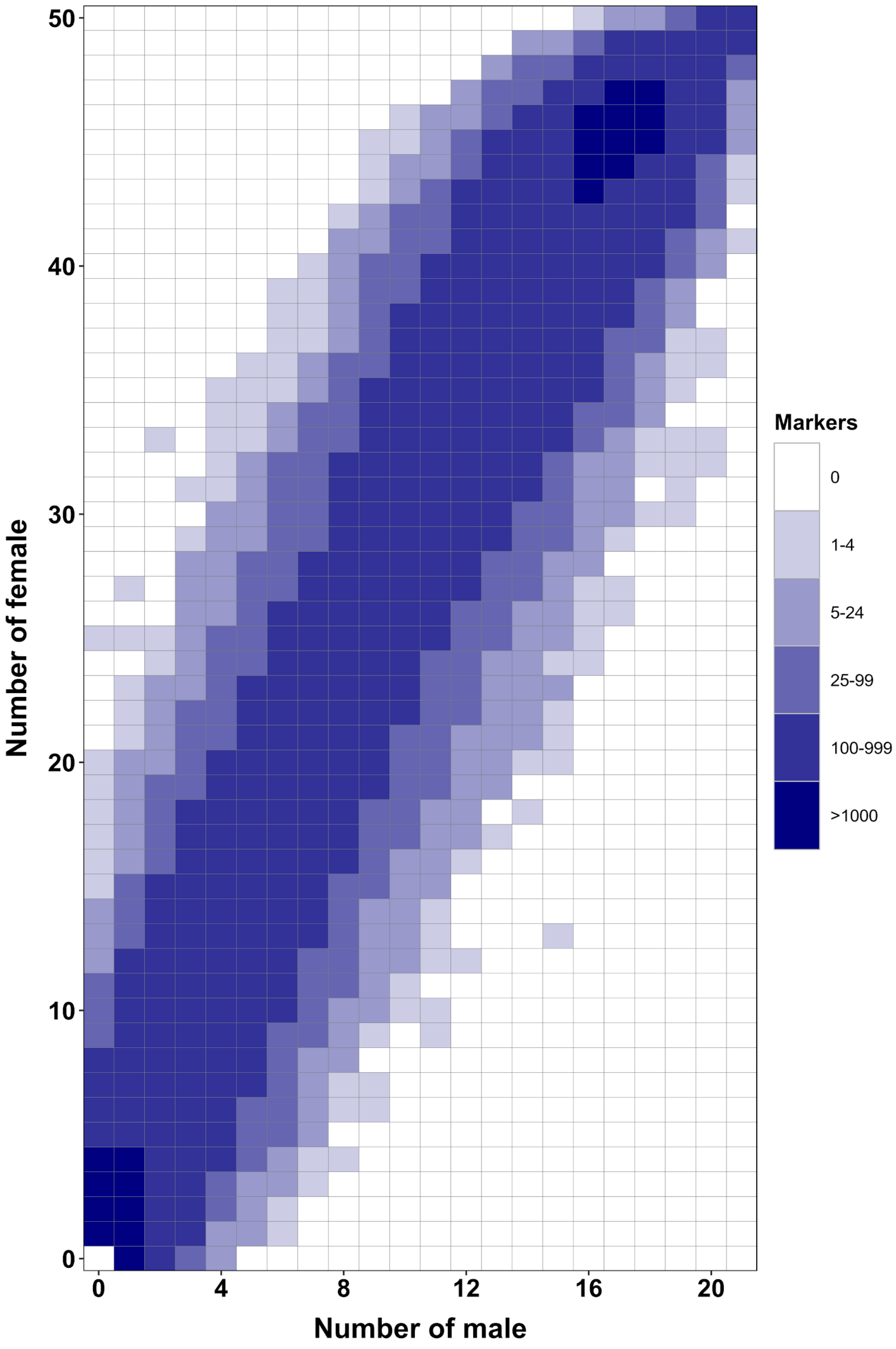


**Figure S7**: **RAD-Seq analysis in a second *Esox masquinongy* population from Quebec (Canada) supports the lack of sex-specific marker in this population.** Each tile plot shows the distribution of non-polymorphic RAD-Seq markers between phenotypic males (horizontal axis) and phenotypic females (vertical axis). The intensity of color for a tile corresponds to the number of markers present in the respective number of males and females. No tiles for which the association with phenotypic sex is significant (Chi-squared test, $p<0.05$ after Bonferroni correction) were detected in this analysis. RAD-Seq reads data and samples information are found under NCBI bioproject PRJNA512459.

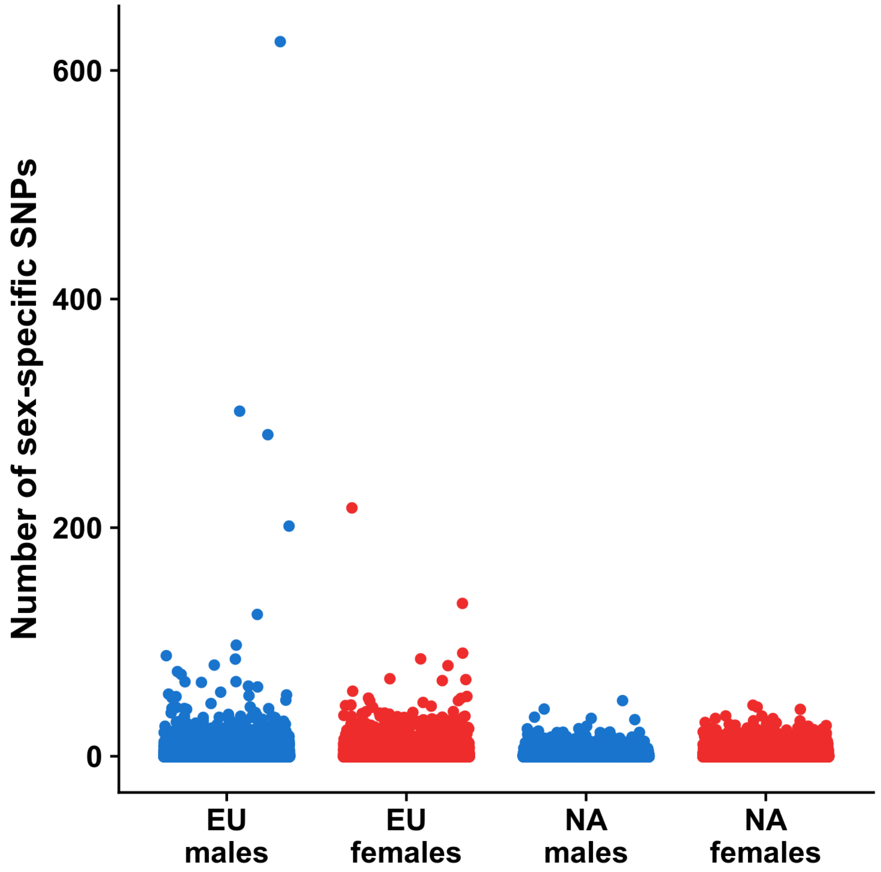


**Figure S7: Number of sex specific SNPs identified for both an European and an North American population**. Sex-specific Pool-Seq reads were mapped to the reference genome of a female *E. lucius* (GCA_004634155.1). Number of male or female specific SNPs in 50Kb non-overlapping windows were computed and plotted for each dataset. Male data is presented in blue and female data is presented in red.

### Supplementary tables

| Metric | Value |
|---|---|
| **Number of scaffolds** | **1231** |
| **Total size of scaffolds** | **919922954** |
| Longest scaffold | 29788268 |
| Shortest scaffold | 517 |
| Number of scaffolds > 1K nt | 1225 (99.5.0%) |
| Number of scaffolds > 10K nt | 1103 (89.6%) |
| Number of scaffolds > 100K nt | 665 (54%) |
| Number of scaffolds > 1M nt | 185 ( 15%) |
| Number of scaffolds > 10M nt | 14 (1.1%) |
| Mean scaffold size | 747,297 |
| Median scaffold size | 127,613 |
| N50 scaffold length | 3,816,572 |
| L50 scaffold count | 51 |
| scaffold %A | 28.98 |
| scaffold %C | 20.98 |
| scaffold %G | 21.03 |
| scaffold %T | 29 |
| scaffold %N | 0 |
| **Complete BUSCOs** | **94.4** |
| Complete single-copy BUSCOs | 90.9 |
| Complete duplicated BUSCOs | 3.50% |
| Fragmented BUSCOs | 2.80% |
| **Missing BUSCOs** | **2.80%** |
| **Complete BUSCOs (ref assembly GCA_004634155.1)** | **4310 (94.1%)** |
| Complete single-copy BUSCOs (ref assembly GCA_004634155.1) | 4142 (90.4%) |
| Complete duplicated BUSCOs (ref assembly GCA_004634155.1) | 168 (3.7%) |
| Fragmented BUSCOs (ref assembly GCA_004634155.1) | 160 (3.5%) |
| **Missing BUSCOs (ref assembly GCA_004634155.1)** | **114 (2.4%)** |

**Table S1**: Assemblathon and BUSCOs metrics for the new genome assembly with additional Nanopore reads of a genetic European male of *E. lucius*.

| Species | Sampling location | Sample collectors | Sexing method | Sampling year | Sample size | Males | Females | Homology cloning | Sex-linkage | RAD-Seq | Pool-Seq | WGS |
|---|---|---|---|---|---|---|---|---|---|---|---|---|
| *Esox lucius* | France | Y.G., Q.P | Gamete production | 2018 | 221 | 161 | 60 | | ✓ | ✓ | ✓ | ✓ |
| *Esox lucius* | Sweden | R.G | Gonad observation | 2012 | 40 | 20 | 20 | | ✓ | | | |
| *Esox lucius* | Poland | K.O | Gamete production | 2013 | 40 | 20 | 20 | | ✓ | | | |
| *Esox lucius* | Xinjiang, China | H.H | Gonad observation | 2018 | 10 | 5 | 5 | | ✓ | | | |
| *Esox lucius* | British Columbia, Canada | B.K., E.R | Gonad observation | 2010-2012 | 20 | 6 | 14 | | ✓ | | | |
| *Esox lucius* | New Jerseys, USA | B.K., E.R | Gamete production | 2011-2012 | 10 | 3 | 7 | | ✓ | | | |
| *Esox lucius* | Manitoba, Canada | B.K., E.R | Gamete production | 2010-2013 | 8 | 3 | 5 | | ✓ | | | |
| *Esox lucius* | Quebec, Canada | L.B | Gamete production | 2009-2010 | 124 | 62 | 62 | | ✓ | ✓ | ✓ | |
| *Esox lucius* | Alaska, USA | J.A.L | Gonad observation | 2018 | 38 | 19 | 19 | | ✓ | | | |
| *Esox aquitanicus* | France | G.P.J.D | External phenotype | 2013 | 3 | 1 | 2 | ✓ | ✓ | | | |
| *Esox cisalpinus* | Italy | G.P.J.D | External phenotype | 2014 | 4 | 2 | 2 | ✓ | ✓ | | | |
| *Esox reichertii* | Heilongjiang,China | S.C | Gonad observation | 2015 | 21 | 11 | 10 | | ✓ | | | |
| *Esox reichertii* | * | * | NR | * | 10 | NA | NA | ✓ | | | | |
| *Esox masquinongy* | Iowa, USA | F.W.G | Gamete production | 2015 | 45 | 27 | 18 | | ✓ | ✓ | ✓ | |
| *Esox masquinongy* | Quebec, Canada | L.B | External phenotype** | 2010-2015 | 40 | 20 | 20 | | ✓ | ✓ | | ✓ |
| *Esox masquinongy* | Wisconsin, USA | W.A.L | Gamete production | 2002-2013 | 122 | 61 | 61 | | ✓ | | | |
| *Esox americanus (americanus)* | Mississippi, USA | E.S., S.S.C | Gonad histology | 2019 | 10 | 6 | 4 | | ✓ | | | |
| *Esox americanus (americanus)* | * | * | NR | * | 10 | NA | NA | ✓ | | | | |
| *Esox americanus (vermiculatus)* | * | * | NR | * | 11 | NA | NA | ✓ | | | | |
| *Esox niger* | Quebec, Canada | L.B | Gonad histology | 2017 | 13 | 5 | 8 | | ✓ | | | ✓ |
| *Novumbra hubbsi* | Washington, USA | K.M.N., R.T., P.D | External phenotype*** | 2010-2012 | 45 | 23 | 22 | | ✓ | ✓ | | ✓ |
| *Dallia pectoralis* | Alaska, USA | J.H.P., F.v.H | Gonad observation | 2014 | 60 | 30 | 30 | | | ✓ | ✓ | ✓ |
| *Umbra pygmaea* | Belgium | H.V., Y.G | Gonad observation | 2013 | 62 | 31 | 31 | | | ✓ | | ✓ |

**Table S2**: Information on the different Esociformes species, sample collectors, sexing method and experiments performed in this study. * = Samples from (Ouellet-Cauchon et al., 2014)[39]. ** Sex was recorded in this *E. masquinongy* population based on the urogenital pores morphology[40]. *** Sex was recorded in *N. hubbsi* based on the specific coloration of males during the breeding season[41]. NR = Phenotypes not recorded. NA = Not applicable (sex phenotypes not recorded). Museum collection numbers are: MNHN 2014-2719, MNHN 2014-2720, MNHN 2014-2721, MNHN 2014-2722, and MNHN 2014-2723 for *E. cisalpinus* and MNHN 2013-1246, MNHN 2013-1245 and MNHN 2013-838 for *E. aquitanicus*.

| Species | Sex | Sequencing approach | Format | Instrument | Total number of reads |
|---|---|---|---|---|---|
| *Esoc lucius (European population)* | male | Pool-Seq | 2*150 bp | NovaSeq S4 | 2.59E+08 |
| *Esox lucius* (European population) | female | Pool-Seq | 2*150 bp | NovaSeq S4 | 2.73E+08 |
| *Esox lucius* (Canadian population) | male | Pool-Seq | 2*150 bp | NovaSeq S4 | 2.81E+08 |
| *Esox lucius* (Canadian population) | female | Pool-Seq | 2*150 bp | NovaSeq S4 | 2.92E+08 |
| *Esox masquinongy* (US population) | male | Pool-Seq | 2*150 bp | NovaSeq S4 | 1.95E+08 |
| *Esox masquinongy* (US populatio*n)* | female | Pool-Seq | 2*150 bp | NovaSeq S4 | 2.52E+08 |
| *Dallia pectoralis* (Alaska) | male | Pool-Seq | 2*150 bp | NovaSeq S4 | 1.99E+08 |
| *Dallia pectoralis* (Alaska) | female | Pool-Seq | 2*150 bp | NovaSeq S4 | 2.80E+08 |
| *Esox lucius* (European population) | male | WGS | 2*250 bp | HIseq2500 | 4.71E+08 |
| *Esox masquinongy* (US population) | male | WGS | 2*250 bp | HIseq2500 | 2.32E+08 |
| *Esox masquinongy (*US population) | female | WGS | 2*250 bp | HIseq2500 | 4.63E+08 |
| *Esox masquinongy (*US population*)* | female | WGS | 2*250 bp | HIseq2500 | 1.68E+08 |
| *Esox niger* (Quebec, canada) | male | WGS | 2*250 bp | HIseq2500 | 2.07E+08 |
| *Novumbra hubbsi* (Washington, USA) | male | WGS | 2*250 bp | HIseq2500 | 2.25E+08 |
| *Dallia pectoralis* (Alaska) | male | WGS | 2*250 bp | HIseq2500 | 2.17E+08 |
| *Umbra pygmaea* (Belgium) | male | WGS | 2*250 bp | HIseq2500 | 4.19E+08 |

**Table S3**: Sequencing information for the Pool-Seq and Whole-genome sequencing (WGS) performed in this study.

| Dataset | Total # of reads | Total # of markers | Average # of markers / individual | Max # of markers | Min # of markers | Min. depth | Retained marker |
|---|---|---|---|---|---|---|---|
| *Dallia pectoralis* | 1.15E+08 | 4.12E+06 | 1.28E+05 | 9.62E+04 | 2.13E+05 | 5 | 4.85E+04 |
| *Esox lucius Canadian* Quebec | 1.33E+08 | 8.44E+06 | 2.13E+05 | 1.36E+05 | 4.35E+05 | 5 | 1.68E+05 |
| *Esox lucius Canadian* British Columbia | 1.12E+08 | 4.53E+06 | 1.81E+05 | 6.08E+04 | 3.34E+05 | 5 | 1.39E+05 |
| *Esox masquinongy* Quebec | 1.46E+08 | 1.29E+07 | 4.29E+05 | 2.34E+05 | 6.34E+05 | 5 | 2.49E+05 |
| *Esox masquinongy* Iowa | 1.71E+08 | 1.57E+07 | 4.05E+05 | 4.04E+04 | 7.33E+05 | 5 | 3.09E+05 |
| *Novumbra hubbsi* | 2.15E+08 | 1.47E+07 | 4.16E+05 | 2.74E+04 | 6.80E+05 | 5 | 1.83E+05 |
| *Umbra pygmaea* | 1.54E+08 | 6.50E+06 | 2.09E+05 | 1.69E+05 | 2.62E+05 | 5 | 4.53E+06 |

**Table S4**: Total number of reads and markers and range of markers among individuals for each RAD-Seq dataset. The number of markers retained, correspond to the number of markers present with depth higher than Min. depth in at least one individual.

| **Assembly** | **Sex** | **Scaffolds** | **Total scaffold size** | **Longest scaffold** | Mean scaffold size | Median scaffold size | N50 (Kb) | L50 | **Complete BUSCOs (%)** | **Single-copy BUSCOs (%)** | **Missing BUSCOs (%)** |
|---|---|---|---|---|---|---|---|---|---|---|---|
| *Dallia pectoralis* | Male | 989,835 | 0.86 G | 190,332 | 864 | 255 | 5.08 | 30,350 | 65.1 | 62.9 | 17 |
| *Esox masquinongy* | Male | 483,942 | 1.09 G | 376,038 | 2260 | 259 | 32.5 | 8,594 | 80 | 77.3 | 8.7 |
| *Esox masquinongy* | Female | 660,189 | 1.14 G | 381,697 | 1728 | 245 | 33.4 | 8,502 | 80.7 | 77.8 | 8.8 |
| *Esox niger* | Male | 331,978 | 0.86 G | 384,029 | 2590 | 248 | 35.1 | 6,261 | 81.7 | 78.6 | 8.3 |
| *Novumbra hubbsi* | Male | 396,170 | 0.76 G | 481,803 | 1909 | 259 | 28.0 | 6,674 | 78.5 | 76.2 | 10.1 |
| *Umbra pygmaea* | Male | 1,757,182 | 1.95G | 202,595 | 1110 | 244 | 8.82 | 47,154 | 64.9 | 62.6 | 17.7 |

**Table S5**: Assemblathon and BUSCOs metrics for draft genome assembly for the Esociformes species.

| Primer name | Sequence | Sequence used for primer design |
|---|---|---|
| SeqAMH1Fw1 | GAAAGACACTGGCTCACAG | *E. lucius amhby* |
| SeqAMH1Fw2 | TGGCACCATCTCTTTCAC | *E. lucius amhby* |
| SeqAMH1Fw3 | TCCAGTCCCGTTGTTCTC | *E. lucius amhby* |
| SeqAMH1Fw4 | CAACATGGTGGCAACTAAGTG | *E. lucius amhby* |
| SeqAMH1Rev1 | TTTCCCTCTGATGGCATTCC | *E. lucius amhby* |
| SeqAMH1Rev2 | CTGTTGGTGGTTCTTTGC | *E. lucius amhby* |
| SeqAMH1Rev3 | TGGCAGTGGAAGATATTAGC | *E. lucius amhby* |
| SeqAMH1Rev4 | GGTAATATTTGTGCCCTGTG | *E. lucius amhby* |
| seqAMH2_Fw1 | TGTTAGATGCAGACGTGAG | *E. lucius amhby* |
| seqAMH2_Rev1 | CACTTTCCTTGTCCCAACC | *E. lucius amhby* |
| seqAMH2_Fw2 | CCATCGTTCCCAACACTTCTC | *E. lucius amhby* |
| seqAMH2_Rev2 | AGGACATCACTGAGGAACC | *E. lucius amhby* |
| seqAMH2_Fw3 | GTAATTGTCTCCATCAGAGCGT | *E. lucius amhby* |
| seqAMH2_Rev3 | CACTCACACTTCGTTGCCAC | *E. lucius amhby* |
| ConserveAMH1_F1 | GTTACTTTTTCTGCCTAGCGTGA | Conserved region on *amhby* orthologs |
| ConserveAMH1_F2 | GTGATAGGCACTGTAACACACCA | Conserved region on *amhby* orthologs |
| ConserveAMH1_R1 | CTATTACTAGTGTGGATAAGGCCG | Conserved region on *amhby* orthologs |
| ConserveAMH1_R2 | CCTCTTCACCCTCAGTGAGC | Conserved region on *amhby* orthologs |
| Con_Amha_F1 | GTCACTATAGGTTGGTTCATC | Conserved region on *amha* orthologs |
| Con_Amha_R1 | CGATTTGCCATTTCAGGTG | Conserved region on *amha* orthologs |
| Con_Amha_F2 | CACGATCTGCAGCCTTACAA | Conserved region on *amha* orthologs |
| Con_Amha_R2 | CTTCAGTAGTAGCAGGGCACG | Conserved region on *amha* orthologs |
| ESAA_Amhby_gapF1 | TAGTAACGGCCTTATCCACAC | *E. americanus amhby* |
| ESAA_Amhby_gapR1 | GGCAGGAGTCATGTATCAACAG | *E. americanus amhby* |
| ESAA_Amhby_spcR1 | GTCACAGCTAAATCAGGTCAC | *E. americanus amhby* |
| Novum_Amha_F1 | GTTTATTCAACTGATGTGGGAC | *N. hubbsi amha* |
| Novum_Amha_R1 | GAAAGCATATCGTCAACAACC | *N. hubbsi amha* |
| Novum_Amhby_F1 | CAAGATGCCAACAAACAGAG | *N. hubbsi amhby* |
| Novum_Amhby_R1 | TATGGATAATAGCCTGAACCTG | *N. hubbsi amhby* |

**Table S6**: Primers used in this study to amplify *amha* and *amhby* sequences from the Esociformes.

| | dN/dS ratio (comparing with *U.pygmaea amh* ) | |
|---|---|---|
| **Species** | ***amha*** | ***amhby*** |
| *Dallia pectoralis* | 0.37 | NA |
| *Novumbra hubbsi* | 0.4301 | truncated |
| *Esox niger* | 0.3732 | truncated |
| *Esox americanus americanus* | 0.3826 | 0.3579 |
| *Esox americanus vermiculatus* | 0.3933 | 0.3561 |
| *Esox masquinongy* | 0.3877 | 0.3808 |
| *Esox reichertii* | 0.3786 | 0.3797 |
| *Esox aquitanicus* | 0.3706 | 0.3848 |
| *Esox cisalpinus* | 0.3629 | 0.3876 |
| *Esox lucius* | 0.3678 | 0.3804 |

**Table S7**: dN/dS ratio between the *amh* paralogs in different Esociformes and *amh* of *U. pygmaea.*

| Model name in PALM | Log-likelihood of the model |
|---|---|
| M0 | -5121.89 |
| M1a | -5123.61 |
| M2 | -5123.61 |
| M7 | -5123.41 |
| M8 | -5123.28 |
| Free-ratio model | -5139.74 |
| Branch-site model | -5122.8 |

**Table S8**: Log-likelihood of different selection models tested on *amha* and *amhby* orthologs of the Esociformes.

| | James Pond | Powell Creek | Stearns Wetland | S. Hanaford Creek | Conner Creek | Steamboat Bog |
|---|---|---|---|---|---|---|
| James Pond | | 0.246 | 0.491 | 0.424 | 0.301 | 0.169 |
| Powell Creek | | | 0.243 | 0.205 | 0.075 | 0.248 |
| Stearns Wetland | | | | 0.033 | 0.25 | 0.567 |
| S. Hanaford Creek | | | | | 0.213 | 0.475 |
| Conner Creek | | | | | | 0.266 |

**Table S9**: Pairwise comparison of Fst values between different populations of *Novumbra hubbsi* used in the RADSex analysis.

| Dataset | Number of *SbfI* cleavage sites | Number of RAD marker/Mb | Genome assembly size |
|---|---|---|---|
| *E. lucius* (Canadian population) | 14,818 | 31.8 | 0.84 G |
| *E. masquinongy* (US population) | 20,852 | 38.4 | 1.09 G |
| *N. hubbsi* | 15,678 | 41.6 | 0.76 G |
| *D. pectoralis* | 13,158 | 30.8 | 0.86 G |
| *U. pygmaea* | 20,227 | 26.2 | 1.54 G |

**Table S10**: Estimated number of *SbfI* cutting sites and RAD-Seq marker frequency estimated for *E. lucius*, *E. masquinongy*, *N. hubbsi*, *D. pectoralis* and *U. pygmaea* based on the size of draft genome assembly.
